# An extracellular complex between CLE-1/collagen XV/XVIII and Punctin/MADD-4 defines cholinergic synapse identity

**DOI:** 10.1101/2025.03.26.645569

**Authors:** Mélissa Cizeron, Anaïs Dumas, Suzanne Le Reun, Laure Granger, Maëlle Jospin, Océane Romatif, Camille Vachon, Delphine Le Guern, Aurore-Cécile Valfort, Jean-Louis Bessereau

## Abstract

The precise localization of postsynaptic receptors opposite neurotransmitter release sites is essential for synaptic function. This alignment relies on adhesion molecules, intracellular scaffolds, and a growing class of extracellular scaffolding proteins. However, how these secreted proteins are retained at synapses remains unclear. We addressed this question using *C. elegans* neuromuscular junctions, where Punctin, a conserved extracellular synaptic organizer, positions postsynaptic receptors. We identified CLE-1, the ortholog of collagens XV/XVIII, as a key stabilizer of Punctin. Punctin and CLE-1B, the main isoform present at neuromuscular junctions, form a complex and rely on each other for synaptic localization. Punctin undergoes cleavage, and in the absence of CLE-1, specific fragments are lost, resulting in the mislocalization of cholinergic receptors to GABAergic synapses. Additionally, CLE-1 regulates receptor levels independently of Punctin. These findings highlight a crucial extracellular complex that maintains synapse identity.

## Introduction

The precise alignment of postsynaptic neurotransmitter receptors with presynaptic release sites is critical for efficient neurotransmission at chemical synapses. Historically, this function has been attributed to adhesion molecules, which form transsynaptic bridges between pre- and postsynaptic domains. However, recent discoveries have revealed a new class of synaptic organizers: extracellular scaffolding proteins^1,2^. Secreted in the extracellular matrix (ECM) by neurons or glial cells, these proteins are now recognized as central players in synapse organization, contributing to synaptogenesis and to the localization of postsynaptic receptors.

Among extracellular scaffolding proteins, cerebellin 1 forms a tripartite complex with the presynaptic adhesion molecule neurexin and the postsynaptic GluD2 receptor which is essential for presynaptic assembly and GluD2 receptor localization^3–5^. While interaction with neurexin provides a mechanism for cerebellin stabilization at synaptic sites, many other extracellular scaffolding proteins lack direct connections to presynaptic adhesion molecules. Thus, the mechanism governing synaptic cleft retention remain unknown for these proteins.

Agrin, which clusters acetylcholine receptors at neuromuscular junctions (NMJs) via LRP4-MuSK signaling^6^, is one of the best characterized extracellular synaptic organizers, yet how it is stabilized at the synaptic cleft is still not fully understood. This question also applies to extracellular scaffolding proteins in the central nervous system, such as neurocan, which induces inhibitory synaptogenesis in cortical neurons^7^, and thrombospondins, a family of five extracellular proteins which induce synaptogenesis through the binding of the α_2_δ-1 subunit of postsynaptic voltage-gated calcium channels^8,9^.

This lack of known presynaptic interactions suggests the involvement of as-yet-unidentified partners that stabilize these extracellular synaptic organizers within the synaptic cleft. To address this, we used *Caenorhabditis elegans*, which allows the precise dissection of synaptic molecular mechanisms with a strong functional conservation^10,11^. We employed an unbiased approach to identify genes essential for the synaptic localization of Punctin, a conserved extracellular scaffolding protein critical for the organization of *C. elegans* NMJs, whose ortholog ADAMTSL3 is required for glutamatergic and GABAergic synapse formation in the murine hippocampus^12^.

Punctin is a member of the ADAMTS-like (ADAMTSL) family, a group of secreted proteins related to the ADAMTS superfamily, but lacking enzymatic activity. Punctin is expressed as long and short isoforms, PunctinL and PunctinS, generated through alternative promoter usage. Structurally, Punctin contains multiple thrombospondin type-1 repeats (TSP), an immunoglobulin (Ig) domain, and a C-terminal PLAC domain. Additionally, PunctinL includes a cysteine-rich domain and a spacer module characteristic of ADAMTS proteins.

In *C. elegans*, NMJs are arranged along two nerve cords and alternate between excitatory (cholinergic) and inhibitory (GABAergic) synapses (Fig. 1A). Muscle cells express acetylcholine and GABA receptors, which are localized opposite the corresponding synaptic terminals by Punctin^13–17^. Punctin localizes three classes of postsynaptic neurotransmitter receptors: homomeric nicotine-sensitive acetylcholine receptors (N-AChRs), heteromeric levamisole-sensitive acetylcholine receptors (L-AChRs), and GABA_A_ receptors (GABARs, Fig. 1A).

**Figure 1:**
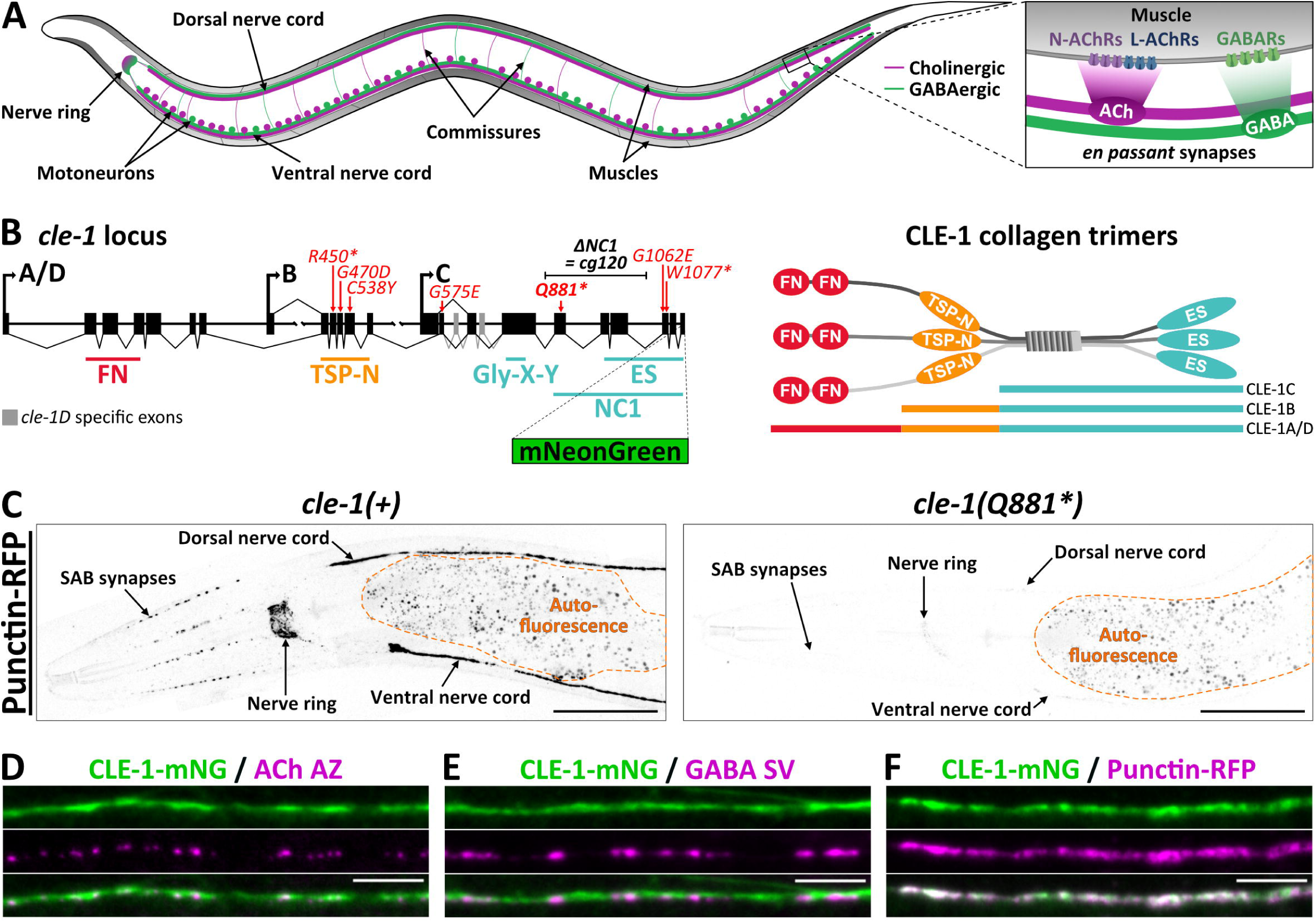
CLE-1/Collagen XV/XVIII localizes Punctin at synapses. A, Schematics of the nematode neuromuscular system and neuromuscular junctions (inset). A subset of motoneurons is represented for simplicity. B, Schematics showing the organization of the *cle-1* locus (left), with the localization of the *cg120* deletion (thereafter referred as *ΔNC1*), the 7 alleles retrieved from the screen (red), and the localization of the mNeonGreen (mNG) C-terminal insertion. Schematics showing the proteic organization of CLE-1 trimers. FN, fibronectin type III domain; TSP-N, Thrombospondin N-terminal-like domain; Gly-X-Y, Collagen domain; NC1, Non-collagenous domain 1; ES, endostatin. C, Confocal images showing the expression of the Punctin-RFP KI (*kr373*) in controls and *cle-1(Q881*)* mutants. D-F, Confocal images showing the colocalization of CLE-1-mNG (*qy22*, green) with presynaptic cholinergic active zones (AZ, P_ACh_::CLA-1-BFP, *krSi145*, magenta in D), GABAergic synaptic vesicles (SV, P_GABA_::SNB-1-BFP, *krIs67*, magenta in E) and Punctin-RFP (magenta in F) in L4 larvae. Scale bars: C, 50 μm; D-F, 5 μm.

The molecular identity of postsynaptic domains is dictated by a code based on the differential secretion of Punctin isoforms. PunctinL, released by cholinergic motoneurons, recruits acetylcholine receptors at cholinergic synapses^13,17^. In contrast, PunctinS, secreted by both cholinergic and GABAergic motoneurons, localizes GABARs through dual mechanisms: promoting their recruitment to GABAergic synapses while simultaneously inhibiting their recruitment at cholinergic synapses by antagonizing PunctinL^13–16^. In the absence of Punctin, postsynaptic receptors are mislocalized to extrasynaptic sites, including muscle arms — membrane extensions of muscle cells projecting toward nerve cords^13^. Although the synaptic localization of Punctin is essential for NMJ organization, the mechanisms responsible for its retention at NMJs remain unknown.

Here, we identify CLE-1, a non-fibrillar collagen and the sole *C. elegans* ortholog of collagens XV/XVIII, as a novel extracellular scaffolding protein required for Punctin localization at synapses. CLE-1B, the principal isoform expressed at *C. elegans* NMJs, forms a molecular complex with Punctin that is necessary for their mutual synaptic localization. Interestingly, Punctin undergoes proteolytic cleavage, with CLE-1B playing a crucial role in localizing specific Punctin fragments. The redistribution of Punctin fragments in the absence of CLE-1 leads to mislocalization of acetylcholine receptors to GABAergic synapses. Additionally, CLE-1 regulates the levels of postsynaptic receptors through a Punctin-independent mechanism. These findings reveal a novel extracellular scaffold that orchestrates synapse identity and regulates synaptic receptor content.

## Results

### CLE-1, the ortholog of collagens XV/XVIII, localizes Punctin at synapses

Punctin is an extracellular matrix protein secreted by motoneurons that localizes neurotransmitter receptors in register with the corresponding presynaptic boutons^13^. To identify the mechanisms involved in the synaptic localization of Punctin, we mutagenized a Punctin-RFP knock-in (KI) strain and screened for mutants with abnormal fluorescence patterns. We retrieved 7 mutations in the *cle-1* locus (Fig. 1B). CLE-1 is the ortholog of collagens XV/XVIII, widely expressed non-fibrillar collagens that also carry glycosaminoglycan chains. All *cle-1* mutations caused a drastic decrease in Punctin-RFP levels, as shown for the *Q881\** allele, which we reproduced by CRISPR/Cas9 gene editing in Punctin-RFP KIs (Fig. 1C). Thus, CLE-1 is necessary for Punctin localization at synapses.

The *cle-1* locus encodes 4 isoforms: 2 long (A and D), 1 medium (B) and 1 short (C), which are generated by specific promoters (Fig. 1B). All CLE-1 isoforms share a collagenous region composed of Gly-X-Y repeats and a C-terminal non-collagenous NC1 domain containing the so-called endostatin domain. In mammals, cleavage of the NC1 domain from collagen XVIII by different proteases can release endostatin, a potent anti-angiogenic factor extensively studied for its anti-tumoral effects^18,19^. Long and medium isoforms also contain a domain homologous to the N-terminal domain of Thrombospondin-1 (TSP-N). Long isoforms additionally contain 2 fibronectin type III domains (Fig. 1B).

### CLE-1B is enriched at cholinergic and GABAergic synapses

We first evaluated the expression of CLE-1 at NMJs using CLE-1-mNG, a KI strain in which all CLE-1 isoforms are labeled with mNeonGreen (mNG) in C-ter^20^ (Fig. 1B). CLE-1-mNG was detected all along the dorsal and ventral nerve cords, where it was enriched in puncta that colocalized with both cholinergic and GABAergic presynaptic markers (Fig. 1D-E). This pattern was similar to that of Punctin, which indeed colocalized with CLE-1 along the nerve cords (Fig. 1F). Thus, CLE-1 is enriched at both cholinergic and GABAergic synapses at *C. elegans* NMJs together with Punctin.

To determine which CLE-1 isoforms were present at NMJs, we generated transcriptional reporters for each isoform using CRISPR/Cas9 gene editing. Specifically, we inserted a sequence at the level of the start codon of each isoform, encoding two nuclear localization signals, the red fluorescent protein wormScarlet (wSC), and a trans-splicing sequence for independent mRNA processing. This revealed very different expression profiles (Fig. 2A), consistent with previous studies^21^. CLE-1A/D were only expressed in a subset of neurons in the head. CLE-1B was expressed in some head neurons and in all motoneurons along the ventral cord. CLE-1C exhibited a broader expression profile, including some head neurons, motoneurons along the ventral cord, the so-called CAN neurons, tail neurons, the gonads, and weak expression in muscle cells. We then generated KI strains for each CLE-1 isoform by using CRISPR/Cas9 to insert the mNG sequence fused to each N-terminus (Fig. 2B). CLE-1A/D were mainly present in the nerve ring. CLE-1B was localized in the nerve ring, along the ventral and dorsal nerve cords (DNC), and at NMJs in the head. CLE-1C was present around the pharynx, gonads, muscles and in the epidermis. Strikingly, CLE-1B had a very similar pattern to Punctin (Fig. 1C) and was highly enriched at the DNC (Fig. 2C). In contrast, little CLE-1C and no CLE-1A/D were detected at the DNC (Fig. 2C). Thus, CLE-1B is expressed by motoneurons and is the main isoform present at GABAergic and cholinergic NMJs.

**Figure 2:**
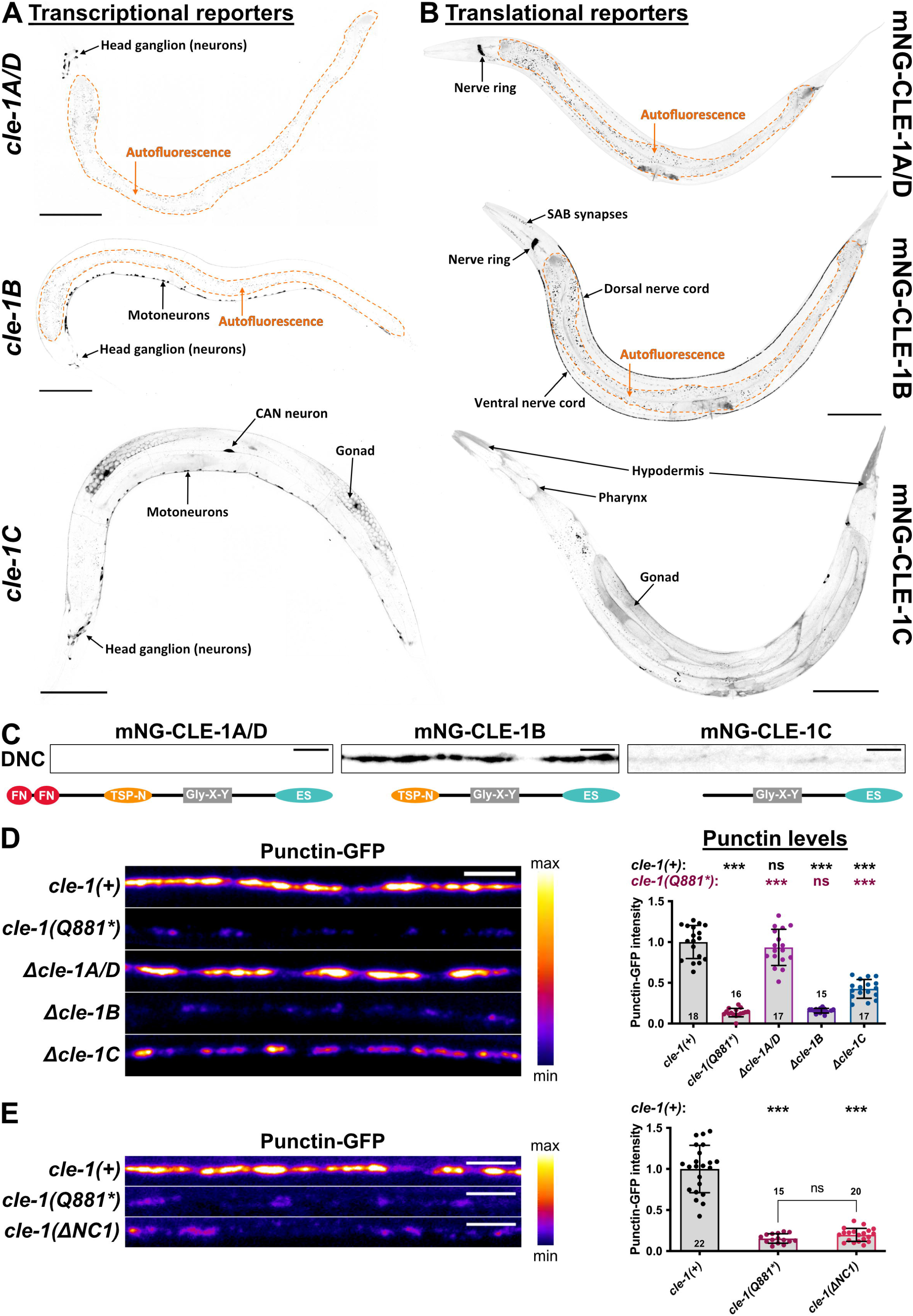
CLE-1B isoform is necessary for Punctin localization at synapses. A-B, Confocal images of transcriptional (A) and translational (B) KI reporters for *cle-1* long (top), medium (middle) and short (bottom) isoforms. C, Confocal images (DNC, dorsal nerve cords in L4 larvae, top) and schematics showing the protein domains (bottom) of CLE-1A/D (left), CLE-1B (middle) and CLE-1C (right). D, Confocal images and quantification of Punctin-GFP intensity at DNCs in controls, *cle-1(Q881*)*, *Δcle-1A/D*, *Δcle-1B* and *Δcle-1C* mutants. Welch’s ANOVA with Dunnett’s T3 multiple comparisons test. E, Confocal images and quantification of Punctin-GFP intensity at DNCs in controls, *cle-1(Q881*)* and *cle-1(ΔNC1)* mutants. Welch’s ANOVA with Dunnett’s T3 multiple comparisons test. Number of animals indicated on graphs. Scale bars: A-B, 100 μm; C-E, 5 μm.

### CLE-1B localizes Punctin at synapses

To test which isoform of CLE-1 was required for the synaptic localization of Punctin, we engineered isoform-specific deletions of *cle-1* in a Punctin-GFP KI strain. Punctin-GFP level was decreased by 85 % at the DNC in *Δcle-1B* mutants, similar to what was observed in *cle-1(Q881*)* mutants (Fig. 2D). Removal of *cle-1A/D* had no effect on Punctin, whereas *cle-1C* deletion decreased Punctin-GFP levels by 57% (Fig. 2D). Since CLE-1C is expressed at low levels at DNCs (Fig. 2C), we wondered whether the decrease of Punctin in *Δcle-1C* mutants might be due to changes in CLE-1B levels. Consistently, introduction of the *Δcle-1C* mutation in the mNG-CLE1B KIs caused a 57% decrease in CLE-1B ((see FigSup 1). It is likely that the decreased Punctin levels observed in *Δcle-1C* mutants result from a decrease in CLE-1B levels. Thus, CLE-1B is the major isoform required for proper Punctin localization at NMJs.

The specific role of the B isoform may suggest that Punctin positioning requires functional domains that are absent in the C isoform, which is also expressed in motoneurons (Fig. 2A). Consistently, our genetic screen identified mutations in the TSP-N domain, which is absent in CLE-1C. However, we observed reduced Punctin levels at synapses with the *Q881\** allele, which could produce truncated versions of CLE-1 lacking the NC1 domain, suggesting that this domain may also be important for Punctin localization (Fig. 1C, 2D-E). Since premature STOP mutations can also cause nonsense-mediated decay of the mRNA, we tested the effect of the *Q881\** allele on CLE-1B expression by introducing this mutation into the mNG-CLE1B KI strain. The mNG-CLE1B signal became undetectable ((see FigSup 2), suggesting that the *Q881\** mutation behaves as a full loss-of-function allele. To investigate the role of the NC1 domain, we therefore introduced in a Punctin-GFP KI strain the *cle-1(cg120)* mutation, referred to as *cle-1(ΔNC1)*, which was previously reported to produce a truncated version of CLE-1 lacking the NC1 domain^21^ (Fig. 1B). Punctin levels were reduced in *cle-1(ΔNC1)* to the same extent as in *cle-1(Q881*)* mutants (Fig. 2E), suggesting that the NC1 domain of CLE-1B is also required for the synaptic localization of Punctin.

### CLE-1 and Punctin mutually localize each other at synapses

Since CLE-1 and Punctin are both ECM proteins that colocalize at NMJs, we wondered whether Punctin might in turn regulate the localization of CLE-1B. We engineered a full deletion of *punctin* in the mNG-CLE-1B KI, because the two loci exhibit genetic linkage. We observed that CLE-1B became completely diffuse at the DNC in *punctin(0)* mutants and its levels were slightly reduced (Fig. 3A). Thus, Punctin is necessary for the synaptic accumulation of CLE-1B.

**Figure 3:**
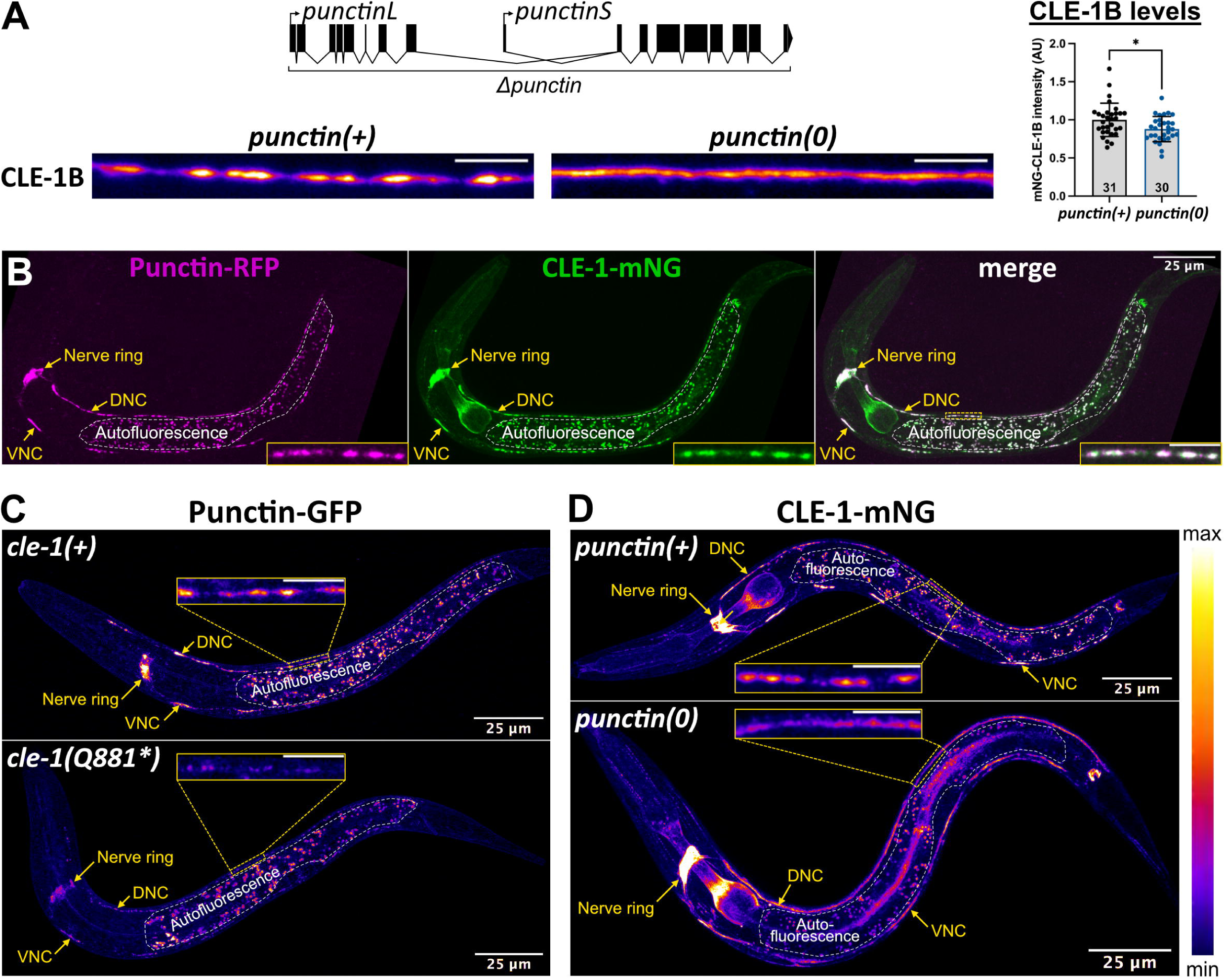
CLE-1 and Punctin localize each other at synapses from early stages. A, Locus descriptions (top), confocal images (bottom) and quantifications (right) of mNG-CLE-1B intensity at DNCs in controls and *punctin(0)* mutants. Mann-Whitney test. B, Confocal images of Punctin-RFP (magenta) and CLE-1-mNG (green) in L1 larvae. C, Confocal images of Punctin-GFP in control and *cle-1(Q881*)* L1 larvae. D, Confocal images of CLE-1-mNG in control and *punctin(0)* L1 larvae. Scale bars: A, 5 μm; B-D, 25 μm, insets: 5 μm.

Since CLE-1B and Punctin require each other for their synaptic localization, we wondered which one might be required first during development. We found that Punctin and CLE-1 were readily detectable and highly colocalized at NMJs at the first larval stage (L1) (Fig. 3B). Remarkably, both Punctin and CLE-1 were mislocalized at the L1 stage in *cle-1(Q881*)* and *punctin(0)* mutants, respectively (Fig. 3C-D). Thus, Punctin and CLE-1 start to localize each other at NMJs early in development.

### CLE-1 and Punctin form a molecular complex

Since CLE-1 and Punctin colocalize at NMJs and functionally interact, we hypothesized that they might be part of the same molecular complex. Immuno-precipitation (IP) of each protein individually retrieved not only the bands corresponding to the predicted isoforms, but also additional species likely corresponding to proteolytic products, which is often observed for ECM proteins.

IP of CLE-1, tagged in C-ter with wSC, retrieved 6 bands (Fig. 4A). The 3 highest bands corresponded to full-length isoforms, as identified by IP of double-tagged isoforms ((see FigSup 3A), and the 3 shorter fragments likely reflected different cleavage site in CLE-1. Interestingly, the smaller band CLE-1(Cter3) could correspond to the release of the endostatin domain, which is cleaved from collagen XVIII in mammals and has specific functions^18,19^.

**Figure 4:**
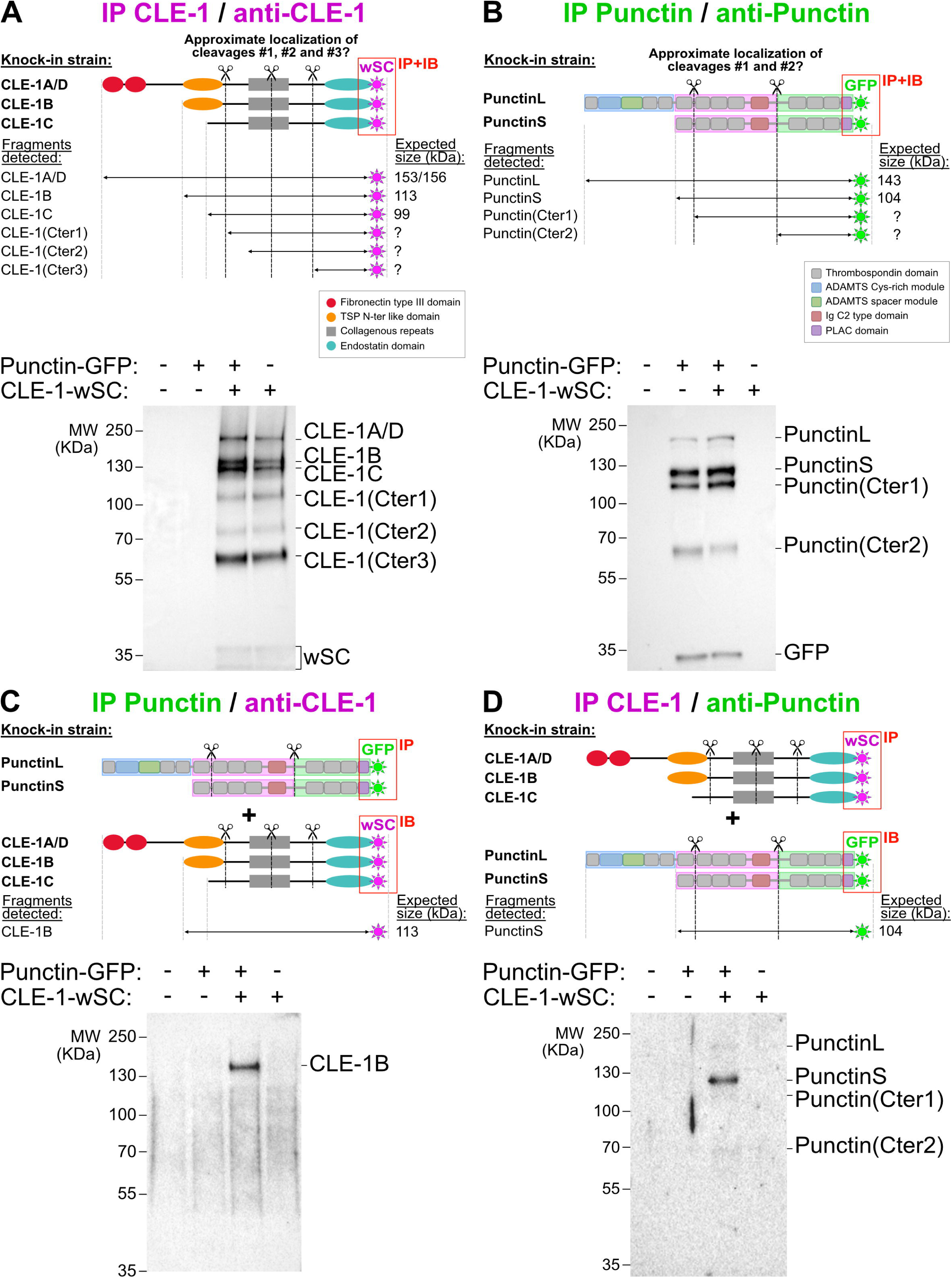
Punctin and CLE-1B form a molecular complex. A-D, Schematics of fusion proteins expressed in KI strains and immunoprecipitation (IP) of CLE-1-wSC (A, D) or Punctin-GFP (B, C), followed by anti-wSC (A, C) or anti-GFP (B, D) immunoblotting (IB) in wild types (lane 1), Punctin-GFP KIs (lane 2), Punctin-GFP + CLE-1-wSC double KIs (lane 3) and CLE-1-wSC KIs (lane 4). Approximate localization of cleavage sites were estimated based on the size of observed bands.

IP of the C-ter tagged Punctin retrieved 4 bands (Fig. 4B), the 2 highest corresponding to full-length PunctinL and PunctinS, as confirmed by IP of double-tagged isoforms ((see FigSup 3B). In addition, we detected 2 lower bands, Punctin(Cter1) and Punctin(Cter2), probably generated by proteolytic cleavage (Fig. 4B). Notably, all bands from CLE-1 and Punctin IPs appeared slightly larger than the predicted sizes (Fig. 4, (see FigSup3), possibly due to glycosylation which is common in ECM proteins.

We then co-immunoprecipitated the two proteins using a double KI strain. Punctin specifically co-immunoprecipitated with the full-length CLE-1B isoform (Fig. 4C). Conversely, CLE-1 IP retrieved mainly PunctinS (Fig. 4D). To test if PunctinL and CLE-1B are nonetheless in the same molecular complex, we specifically immunoprecipitated PunctinL and were able to retrieve CLE-1B ((see FigSup 4). Taken together, these results demonstrate that Punctin and CLE-1B are in the same molecular complex and suggest that this complex may be required for their mutual synaptic localization.

### Punctin fragments localize to specific synapses

We originally proposed that the repertoire of Punctin isoforms present at NMJs, *i. e.* PunctinL and/or PunctinS, controls the cholinergic or GABAergic identity of postsynaptic domains^13^. However, the generation of proteolytic products significantly increases the potential diversity of Punctin species at each synapse. Based on the size of the two Punctin(Cter) fragments, we hypothesized that these bands could result from cleavage at two different sites localized between the TSP4 and TSP5 domains (cleavage #1) and between the Ig and TSP8 domains (cleavage #2), which are predicted to be unstructured based on AlphaFold predictions (Fig. 5A)^22–24^.

**Figure 5:**
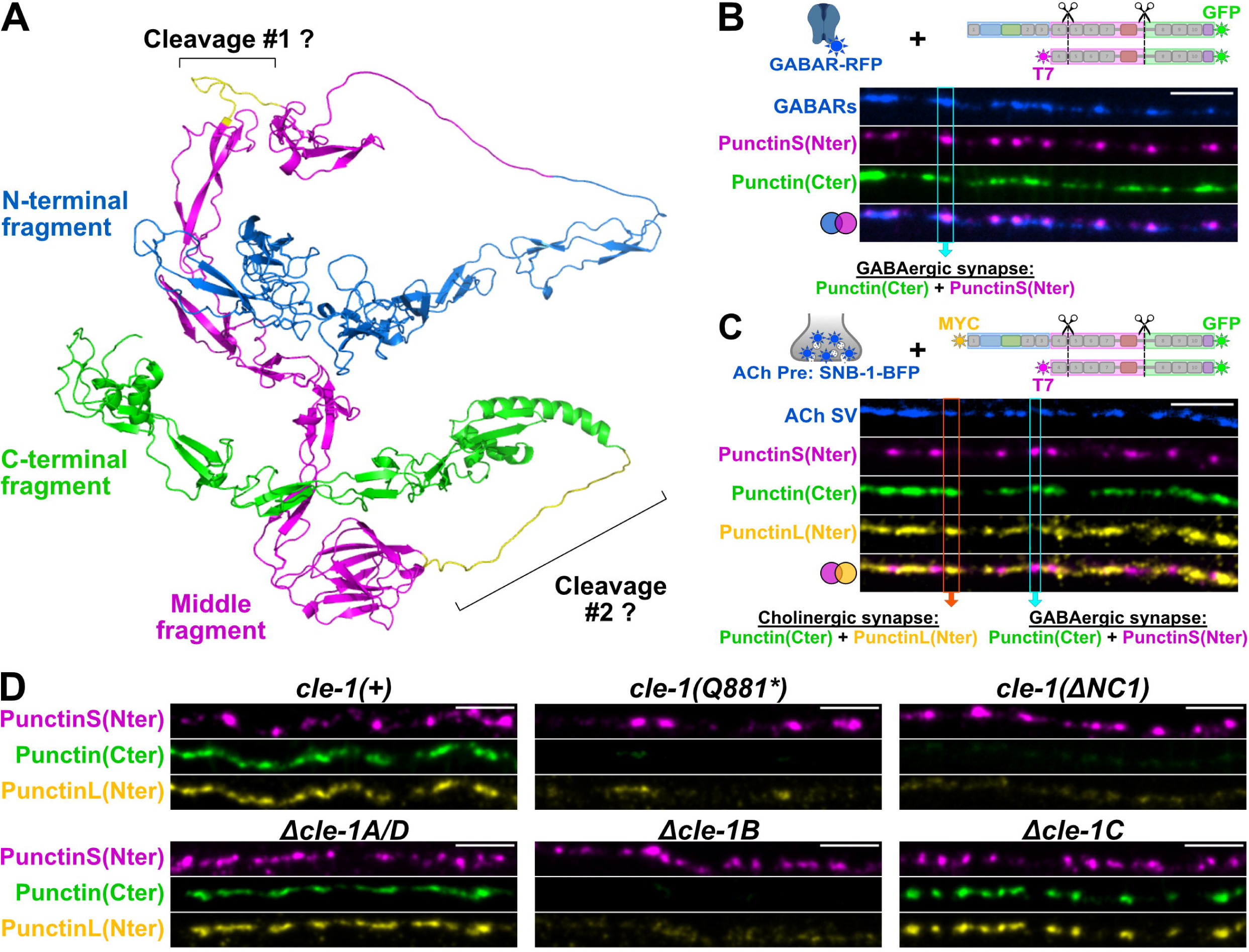
Punctin cleavage generates diverse fragments with specific synaptic distributions. A, Cartoon representation of PunctinL as predicted by AlphaFold. The N-terminal domains are represented in blue (from TSP1 to TSP3), middle domains are magenta (from TSP4 to the IG domain) and C-terminal domains are green (from TSP8 to PLAC), as shown in B. B, Confocal images of GABAR-RFP (blue, *kr296*), Punctin(Cter) (Punctin-GFP, green) and immunofluorescent staining of PunctinS(Nter) (magenta) at the DNC. C, Confocal images of cholinergic boutons (blue, P_ACh_::SNB-1-BFP, *krIs75*), Punctin(Cter) (Punctin-GFP, green) and immunofluorescent staining of PunctinL(Nter) (yellow) at the DNC. D, Confocal images of Punctin(Cter) (Punctin-GFP, green) and immunofluorescent staining of PunctinL(Nter) (yellow) and PunctinS(Nter) (magenta) at the DNC of controls (*kr421*, MYC-PunctinL-GFP+T7-PunctinS-GFP), *cle-1(Q881*)*, *cle-1(ΔNC1)*, *Δcle-1A/D*, *Δcle-1B* and *Δcle-1C* mutants. Scale bars: 5 μm.

We analyzed the localization of each Punctin fragment using immunostaining of a Punctin triple-tagged KI strain, where PunctinL(Nter), PunctinS(Nter) and Punctin(Cter) are tagged with 3xMYC, T7 and GFP, respectively (Fig. 5B, C). Surprisingly, PunctinS(Nter) localized specifically at GABAergic synapses and was absent from cholinergic synapses (Fig. 5B, C), even though PunctinS is expressed in both cholinergic and GABergic motoneurons^13,25^. In contrast, Punctin(Cter), which labels both long and short Punctin isoforms, was present at both cholinergic and GABAergic synapses, as expected (Fig. 5B, C). Finally, PunctinL(Nter), which is expressed only by cholinergic motoneurons^13,25^, was present solely at cholinergic synapses (Fig. 5C). The absence of PunctinL(Nter) at GABAergic synapses was confirmed by its lack of colocalization with PunctinS(Nter) (Fig. 5C). Thus, cholinergic NMJs contain fragments that include Punctin(Cter) and PunctinL(Nter) extremities, but not the PunctinS(Nter), while GABAergic synapses comprise fragments that include Punctin(Cter) or PunctinS(Nter) termini.

### CLE-1B differentially localizes specific Punctin fragments

Since our genetic screen was performed with a C-ter-tagged Punctin, we wondered whether CLE-1 might have distinct effects on the different Punctin fragments. First, we showed that Punctin cleavage was not altered by the absence of CLE-1 in *cle-1(Q881*)* mutants ((see FigSup 5). We then examined the distribution of Punctin extremities in *cle-1* mutants. Strikingly, the localization of PunctinS(Nter) at GABAergic NMJs was unchanged in *cle-1(Q881*)* and *cle-1(ΔNC1)* mutants as compared to the wild type, whereas both PunctinL(Nter) and Punctin(Cter) levels were severely affected (Fig. 5D). Specific deletion of the long isoforms *cle-1A/D* or the short isoform *cle-1C* did not affect the pattern of any Punctin fragment, while deletion of *cle-1B* recapitulated the phenotype seen in *cle-1(Q881*)* mutants (Fig. 5D).

Therefore, the NC1 domain of CLE-1B is specifically required for the localization of PunctinL(Nter)- and Punctin(Cter)-containing fragments, whereas it is dispensable for the localization of PunctinS(Nter)-containing fragments.

### CLE-1 maintains synaptic identity

Since CLE-1 affects differentially Punctin fragments, this prompted us to examine postsynaptic receptors in *cle-1* mutants. Loss of CLE-1 had a dramatic impact on N-AChRs: in *cle-1(Q881*)* mutants, N-AChR levels were reduced by more than 55 % at DNCs (Fig. 6A) and the remaining receptors were ectopically concentrated at GABAergic synapses (Fig. 6A, B). The level of L-AChR was only slightly decreased (-16 %) ((see FigSup 6A), but they also tended to relocalize to GABAergic synapses ((see FigSup 6B). In contrast, GABARs remained properly localized in register with GABAergic active zones, although their levels were slightly reduced (-27 %) ((see FigSup 6C). Thus, the 3 types of receptors are localized at GABAergic synapses in *cle-1* mutants, highlighting an altered synaptic identity at NMJs.

**Figure 6:**
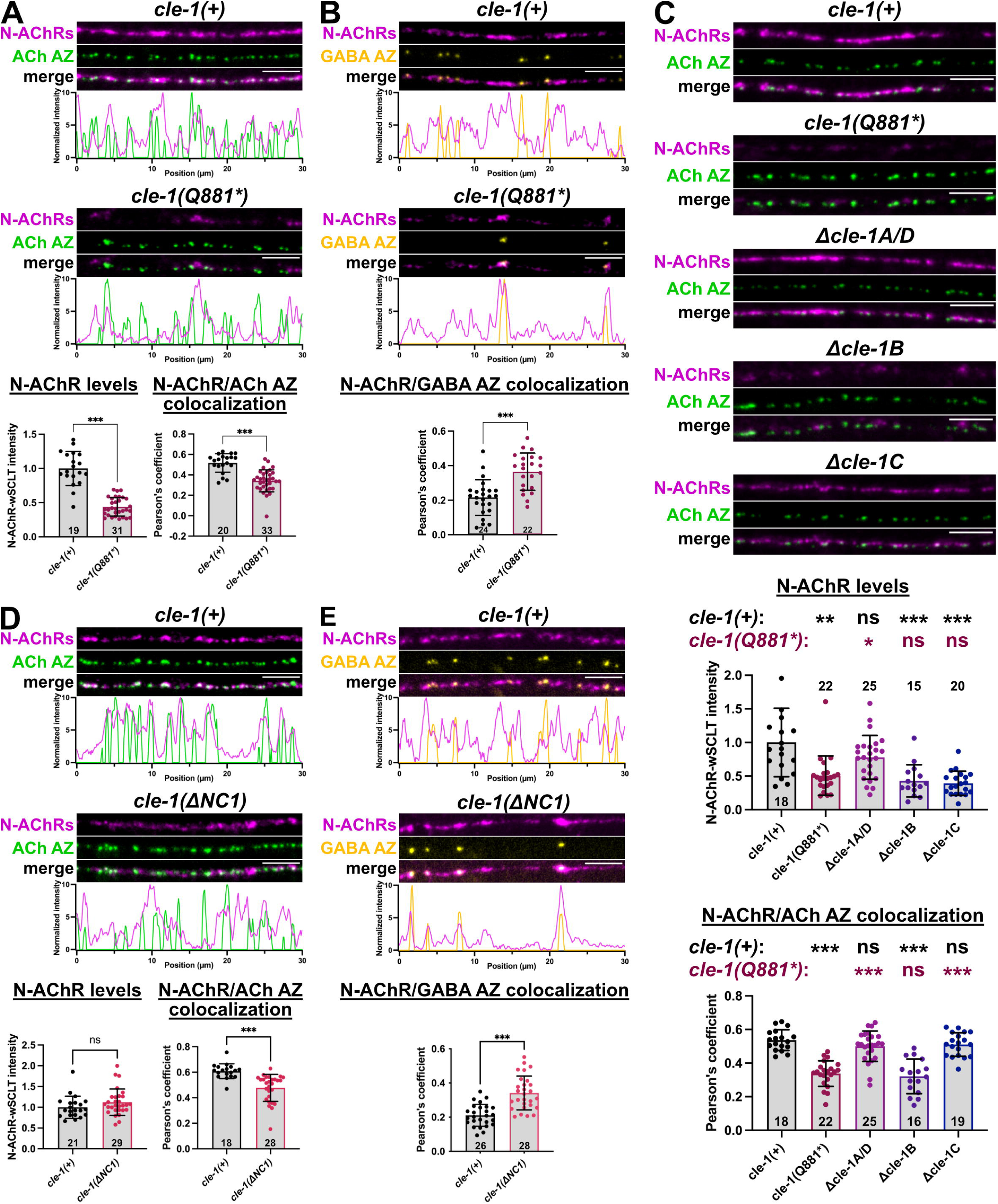
Specific Punctin fragments are redistributed and N-AChRs are relocalized to GABAergic synapses in the absence of CLE-1B. A, Confocal images, fluorescent profiles, quantifications of N-AChR-wSC intensity (magenta, Welch’s t test) and colocalization between N-AChR-wSC and presynaptic cholinergic active zones (AZs, P_ACh_::CLA-1-BFP, *krSi145*, green, Mann-Whitney’s test) in controls and *cle-1(Q881*)* mutants. B, Confocal images, fluorescent profiles and colocalization between N-AChR-wSC and presynaptic GABAergic AZs (P_GABA_::CLA-1-tagBFP, *krSi141*, yellow) in controls and *cle-1(Q881*)* mutants (Student’s t test). C, Confocal images, quantifications of N-AChR-wSC intensity (magenta, Kruskal-Wallis with Dunn’s multiple comparisons tests) and colocalization between N-AChR-wSC and presynaptic cholinergic AZs (green, One way ANOVA with Tukey’s multiple comparisons tests) in controls, *cle-1(Q881*)*, *Δcle-1A/D*, *Δcle-1B* and *Δcle-1C* mutants. D, Confocal images, fluorescent profiles, quantifications of N-AChR-wSC intensity (magenta) and colocalization between N-AChR-wSC and presynaptic cholinergic AZs (green) in controls and *cle-1(ΔNC1)* mutants. Mann-Whitney’s tests. E, Confocal images, fluorescent profiles and colocalization between N-AChR-wSC and presynaptic GABAergic AZs (yellow) in controls and *cle-1(ΔNC1)* mutants (Welch’s t test). Scale bars: 5 μm.

The decrease in N-AChR levels and their relocalization to GABAergic synapses was also observed after specific deletion of *cle-1B* (Fig. 6C). Deletion of *cle-1C* reduced N-AChR levels by 60% without affecting their localization (Fig. 6C). It is possible that the reduced levels of CLE-1B in *Δcle-1C* mutants ((see FigSup 1) are responsible for the decreased N-AChR levels, while the remaining CLE-1B is sufficient to maintain their cholinergic localization. Finally, deletion of *cle-1A* had no effect on N-AChRs (Fig. 6C). Thus, CLE-1B controls both Punctin fragments and postsynaptic receptor localization.

Since the NC1 domain of CLE-1B is required for the localization of Punctin(Cter) and PunctL(Nter) at synapses (Fig. 5D), we assessed its role on postsynaptic receptors. In contrast to what we observed in *cle-1(Q881*)* mutants, we did not see any change in N-AChR or GABAR levels in *cle-1(ΔNC1)* mutants (Fig. 6D, (see FigSup 7). However, N-AChRs relocalized to GABAergic synapses in these mutants (Fig. 6D, E). This suggests a functional dissociation of CLE-1B domains, with the NC1 domain required for cholinergic localization of AChRs, while more N-terminal domains control postsynaptic receptor levels.

Taken together, these data demonstrate that the interplay between CLE-1 and Punctin is critical for maintaining the identity of cholinergic NMJs and tuning the levels of synaptic receptors.

### Loss of CLE-1 causes Punctin-dependent redistribution of SDN-1/Syndecan and N-AChRs to GABAergic synapses

We have previously shown that N-AChR clustering depends on the heparan sulfate proteoglycan SDN-1/syndecan, whose synaptic enrichment requires Punctin^17^. Therefore, we hypothesized that the loss of Punctin fragments at cholinergic synapses in *cle-1* mutants might cause a mislocalization of SDN-1, and subsequently, the abnormal localization of N-AChRs. In the wild type, SDN-1 is present at both cholinergic and GABAergic synapses^17^. In *cle-1* mutants, SDN-1 levels were reduced by approximately 40% and, strikingly, SDN-1 appeared to be much more concentrated at GABAergic synapses than at cholinergic synapses (Fig. 7A, FigSup8). Consistent with previously published results, SDN-1 was further reduced in *punctin(0)* and was no longer enriched at GABA synapses (Fig. 7A). *cle-1(Q881*) punctin(0)* double mutants were similar to *punctin(0)* single mutants, suggesting that the redistribution of Punctin fragments in *cle-1* mutants was responsible for the relocalization of SDN-1 and the subsequent relocalization of N-AChRs (Fig. 7A).

**Figure 7:**
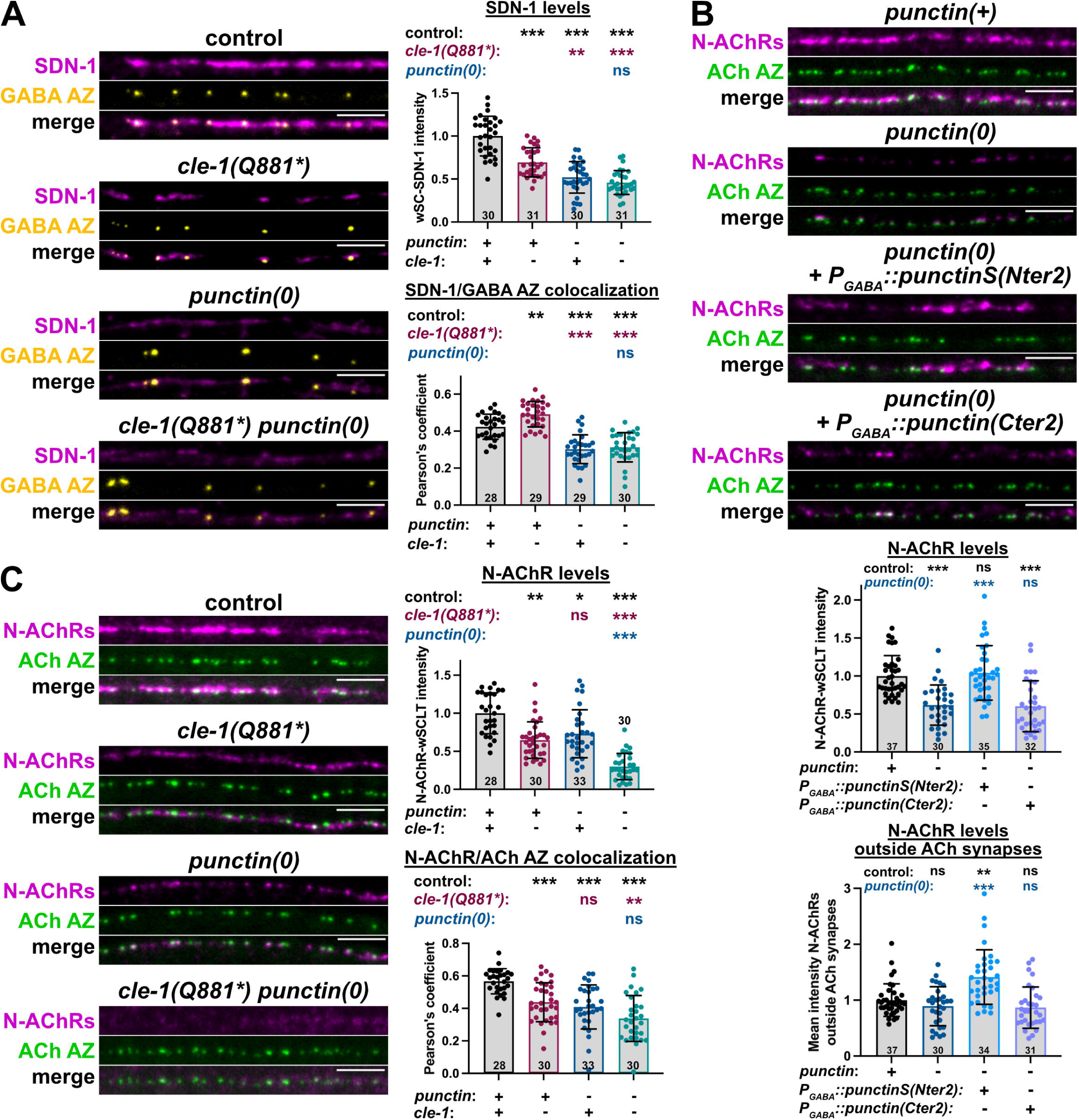
CLE-1 controls N-AChR localization and levels through two parallel pathways. A, Confocal images, quantifications of wSC-SDN-1 intensity (magenta, Welch’s ANOVA with Dunett’s T3 multiple comparison tests) and colocalization between wSC-SDN-1 and GABAergic AZs (yellow, One way ANOVA with Tukey’s multiple comparisons tests) in controls, *cle-1(Q881*)*, *punctin(0)* and *cle-1(Q881*) punctin(0)* mutants. B, Confocal images of N-AChR-wSC (magenta) and cholinergic AZs (green) in control and *punctin(0)* mutants without or with expression of the PunctinS(Nter2) or Punctin(Cter2) fragment in GABAergic neurons. Quantifications of N-AChR-wSC intensity at the cord (One way ANOVA with Tukey’s multiple comparisons tests) and outside cholinergic synapses (Kruskall-Wallis with Dunn’s multiple comparisons tests). C, Confocal images, quantifications of N-AChR-wSC intensity (magenta, Kruskall-Wallis with Dunn’s multiple comparisons tests) and colocalization between N-AChR-wSC and cholinergic AZs (green, One way ANOVA with Tukey’s multiple comparisons tests) in controls, *cle-1(Q881*)*, *punctin(0)* and *cle-1(Q881*) punctin(0)* mutants. Scale bars: 5 μm.

To test this hypothesis, we specifically expressed the N-terminal fragment of PunctinS resulting from cleavage #2, PunctinS(Nter2), which remains at GABAergic synapses in *cle-1* mutants (Fig. 5D), in the GABAergic motoneurons of *punctin(0)* mutants. Expression of PunctinS(Nter2) phenocopied the *cle-1(Q881*)* phenotype, generating large N-AChR clusters at GABAergic synapses (Fig. 7B). In contrary, expression of the C-terminal part of Punctin resulting from cleavage #2, Punctin(Cter2), had no effect on the levels nor localization of N-AChRs (Fig. 7B). These findings suggest that different Punctin fragments have distinct functions. A simple model would therefore predict that the control of N-AChRs is entirely mediated by Punctin. However, when we quantified the levels of N-AChRs remaining at the dorsal nerve cord in *punctin(0)* and *cle-1(Q881*)* single and double mutants, we saw that N-AChRs were further reduced in the double mutants as compared to *punctin(0)* single mutants (Fig. 7C). These results suggest that CLE-1 may also regulate N-AChR levels through a Punctin-independent pathway.

## Discussion

Through an unbiased genetic screen, we identified CLE-1, the ortholog of non-fibrillar collagens XV/XVIII, as a novel synaptic organizer crucial for localizing Punctin, the central extracellular organizer of NMJs^13,15–17^. CLE-1B, the main isoform present at NMJs, localizes specific Punctin fragments generated by proteolytic cleavage. In the absence of CLE-1, synaptic identity is altered, with acetylcholine receptors redistributed to GABAergic synapses. Additionally, CLE-1 regulates postsynaptic receptor levels independently from Punctin. Together, these findings uncover a previously uncharacterized extracellular molecular complex that governs synaptic identity.

### CLE-1, the ortholog of collagens XV/XVIII, organizes *C. elegans* NMJs

CLE-1 shapes ECM composition by positioning Punctin at synapses, thereby dictating synapse identity. CLE-1B and Punctin form a complex and are mutually required for their synaptic enrichment, suggesting that their synaptic retention depends on macromolecular assembly. A similar interaction occurs in *D. melanogaster* between the collagen IV-like Pericardin (Prc) and the ADAMTSL protein Lonely heart (Loh), which mediate adhesion between pericardial cells and cardiomyocytes^26^. However, Prc localization depends unidirectionally on Loh, unlike the reciprocal stabilization of CLE-1B and Punctin, highlighting mechanistic differences^26^.

Punctin undergoes proteolytic cleavage, and specific fragments are lost from the synaptic cleft in the absence of CLE-1B. In *cle-1* mutants, only PunctinS(Nter) termini remain at GABAergic synapses, leading to an enrichment of the transmembrane heparan sulfate SDN-1/Syndecan, whose localization depends on Punctin^17^. SDN-1 redistribution likely drives N-AChR mislocalization, as expression of a chimeric protein with SDN-1 intracellular domains at GABAergic synapses is sufficient to attract them^17^. Thus, CLE-1 directs Punctin fragment localization, positioning SDN-1 to regulate N-AChR localization at cholinergic synapses (Fig. 8).

**Figure 8:**
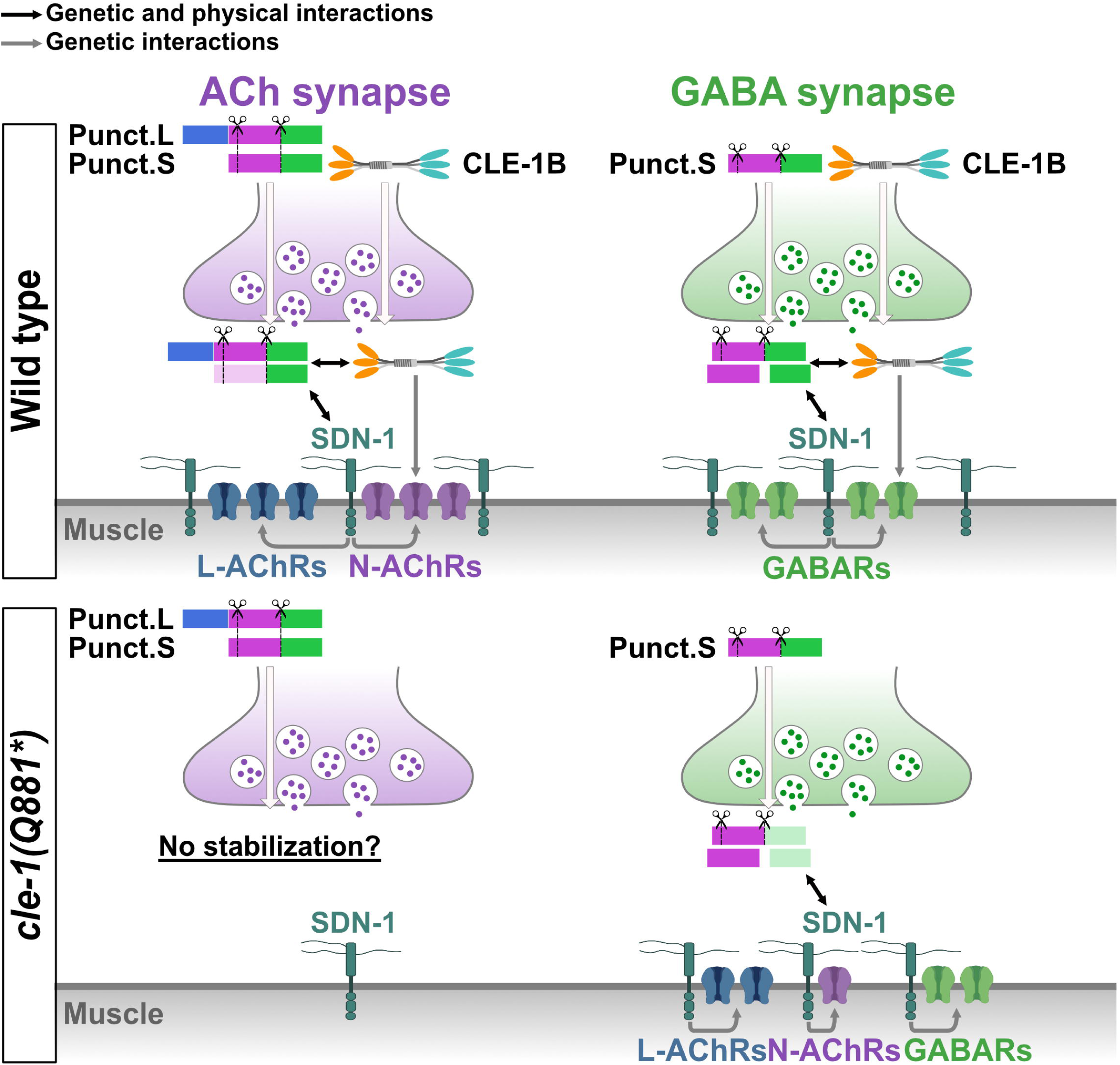
Working model for synapse organization by CLE-1B. See discussion for details. Only a subset of Punctin fragments are represented for simplicity. Transparent fragments indicate regions of the protein that were not observed.

In *cle-1* mutants, L-AChRs are also mislocalized to GABAergic synapses, indicating a broader alteration of synapse identity, while GABARs remain correctly positioned (Fig. 8). The Punctin–CLE-1B complex is therefore crucial for cholinergic identity, and CLE-1 additionally regulates postsynaptic receptor levels through a Punctin-independent mechanism.

This study identifies CLE-1 as a key organizer of postsynaptic structure, adding to its known role in presynaptic assembly. Indeed, CLE-1 constrains GABAergic bouton growth and regulates their number and morphology^27–29^. Thus, CLE-1 is a central NMJ organizer, influencing presynaptic elements, postsynaptic structures, and synaptic cleft composition.

### Full-length CLE-1 is required for Punctin localization

CLE-1, the ortholog of collagens XV/XVIII, belongs to the multiplexin family of collagens, characterized by interrupted triple-helix domains and heparan/chondroitin sulfate chains. In mice, *Col18a1* encodes three collagen XVIII isoforms, while *Col15a1* encodes a single collagen XV isoform, and both are widely expressed^19,30^. CLE-1B shares a similar domain organization with collagen XV and the short isoform of collagen XVIII, comprising a TSP-N domain, collagenous repeats, and a C-terminal non-collagenous NC1 domain, which contains a trimerization region, a hinge with proteolytic cleavage sites, and the endostatin domain.

Endostatin, released by collagen XVIII cleavage, has anti-angiogenic properties with therapeutic potential in cancer^31,32^. However, its precise mechanism of action remains unclear, owing to its pleiotropic effects and its diverse interactions with multiple partners^19^. Endostatin can also have opposite functions depending on its structural form: trimeric NC1 promotes cell migration, whereas cleaved monomeric endostatin inhibits it^33^. In *C. elegans*, *cle-1(ΔNC1)* mutants display cell migration defects rescued by trimeric NC1 but not monomeric endostatin, while endostatin overexpression inhibits migration in wild types, highlighting the antagonistic roles of full-length CLE-1 and monomeric endostatin^21^.

Endostatin also functions at synapses. In *D. melanogaster*, Multiplexin, the ortholog of CLE-1, is present at NMJs where endostatin acts as a trans-synaptic signal to increase presynaptic release when postsynaptic glutamatergic receptors function is reduced, thus maintaining a constant synaptic transmission^34,35^. In mice, collagen XVIII-derived endostatin supports synapse formation between Purkinje cells and climbing fibers *via* α3β1 integrin signaling^36^.

Our data show that the NC1 domain is critical for the localization of Punctin(Cter)- and PunctinL(Nter)-containing fragments. However, Punctin immunoprecipitation retrieved full-length CLE-1B, but not its C-terminal cleavage products, suggesting that monomeric endostatin cannot bind Punctin. While motoneurons express CLE-1B and CLE-1C, only CLE-1B interacts with Punctin, suggesting that the TSP-N domain is important for Punctin binding. Genetic screening supports this hypothesis, identifying three mutations in the TSP-N domain, including two missense mutations. Altogether, these findings suggest that full-length CLE-1B is necessary for proper localization of Punctin fragments, likely through TSP-N-mediated interactions.

### CLE-1 controls postsynaptic receptor levels independently from Punctin

Beyond synaptic identity, CLE-1 regulates neurotransmitter receptor levels at NMJs. In *cle-1(Q881\**) mutants, N-AChR and GABAR levels decrease, whereas *cle-1(ΔNC1)* mutants do not show this effect, suggesting that the endostatin/NC1 domain are dispensable for receptor level maintenance.

Moreover, *cle-1 punctin* double mutants exhibit a greater N-AChR reduction than *punctin* mutants, suggesting that CLE-1 regulates N-AChR levels independently of Punctin. Consistently, SDN-1 levels, which rely entirely on Punctin^17^, are similar between *punctin* mutants and *punctin cle-1* double mutants. Thus, SDN-1 acts downstream of Punctin to regulate N-AChR localization, while CLE-1 influences receptor levels *via* a parallel pathway involving unidentified partners. The netrin receptor UNC-40/DCC regulates GABAR and N-AChR synaptic content and binds Punctin in *C. elegans*^16,17^. In mammals, Cbln4/Cerebellin 4 interacts with DCC^37,38^, demonstrating that DCC can interact with several extracellular synapse organizers, in addition to its canonical netrin ligand. This suggests Punctin and CLE-1B could converge on UNC-40/DCC to regulate N-AChR levels through parallel pathways.

Notably, this study reveals a CLE-1 function independent of its NC1 domain, while most collagen XV/XVIII roles are attributed to their endostatin domain. While functional insights for other domains remain relatively scarce, the TSP-N domain was recently shown to regulate kidney branching morphology in mice^39^. In *D. melanogaster*, the TSP-N domain of Multiplexin controls presynaptic CaV1.2 channel abundance at NMJs^34^, hinting at a conserved role in the regulation of synaptic channel levels. Alternatively, the collagenous repeats of CLE-1B may contribute, as some receptors, like PTPRσ/δ, directly interact with triple-helical collagen chains^40^. Future studies should determine whether the TSP-N domain or collagenous repeats mediate receptor regulation at *C. elegans* NMJs, shedding light on these understudied multiplexin domains.

### Punctin cleavage increases synaptic molecular diversity

Proteolytic cleavage is a key regulatory mechanism for ECM proteins, generating biologically active fragments and increasing molecular diversity^18,41^. This process is critical for synaptic organization, as seen for endostatin release from Multiplexin, which is required for presynaptic homeostasis in *D. melanogaster*^34^, and for neurocan cleavage, where its C-terminal fragment promotes inhibitory synapse formation in mice^7^.

We show that Punctin contains two cleavage sites, possibly increasing its molecular diversity to over ten species. We show that Punctin fragments localize to specific synapses : Punctin(Cter) is present at both types of synapses, PunctinL(Nter) is only at cholinergic synapses and PunctinS(Nter) is only at GABAergic synapses. Moreover, Punctin fragments have different functions, as seen for the PunctinS(Nter2) fragment, which is sufficient to attract N-AChRs at GABAergic synapses, unlike the Punctin(Cter2) fragment. Further studies will be required to determine the function of Punctin cleavage and of the subsequently generated fragments.

PunctinL shares a domain organization with its mammalian orthologs ADAMTSL1 and ADAMTSL3, where putative cleavage events of unknown function have been reported^42,43^. ADAMTSL1 is mainly expressed in skeletal muscle, while ADAMTSL3 is broadly expressed^42,43^, including in the brain where it plays a key role in synaptogenesis and maintenance of inhibitory transmission^12^. Given the conserved role of Punctin and ADAMTSL3 as extracellular synaptic scaffolding proteins, it will be essential to explore whether ADAMTSL3 cleavage contributes to its synaptic functions. More broadly, this study suggests that ADAMTSL protein fragments may possess distinct biological activities compared to the full-length protein. ADAMTSL proteins have pleiotropic functions, including the regulation of fibrillin microfibril formation and TGFβ signaling^44^, opening new avenues for exploring the role of ADAMTSL cleavage in these functions.

### Disease relevance of CLE-1B/Punctin interaction

Elucidating the roles of CLE-1B and Punctin at synapses has substantial medical relevance, as their orthologs are implicated in pathologies. Variants of the *punctin* ortholog *ADAMTSL3* have been identified as a risk factor for schizophrenia^45^ and mutations in *COL18A1*, which encodes collagen XVIII in humans, cause the Knobloch syndrome, a heterogeneous disorder primarily characterized by severe ocular abnormalities that often result in early blindness^19^. Beyond ocular defects, the Knobloch syndrome can include symptoms such as encephalocele or meningocele, with some patients exhibiting autistic traits^46^ or epilepsy^47,48^, potentially reflecting synaptic dysfunction. Additionally, endostatin accumulates at Aβ plaques in Alzheimer’s disease^49,50^, which is characterized by synaptic and neuronal loss.

Functional or physical interactions between ADAMTSLs and collagens XV/XVIII may also contribute to physiopathology. Mutations in *ADAMTSL1*, another *punctin* ortholog, are associated with retinal anomalies, including myopia^51,52^, a hallmark feature of the Knobloch syndrome. This raises the intriguing possibility that *ADAMTSL1* could contribute to retinal symptoms in Knobloch syndrome patients. Furthermore, both *COL18A1* and *ADAMTSL3* exhibit increased expression in optic nerve head astrocytes from patients with primary open-angle glaucoma^53^. Thus, understanding ADAMTSL -collagens XV/XVIII interactions may provide insights to various neurological diseases.

## Acknowledgments

We thank members of the Bessereau lab for providing feedback and Driss Laabid for technical assistance. We are grateful to Lukas Lange for his help during his internship. We thank Fekrije Selimi and Cécile Charrier for critical reading of the manuscript. We acknowledge the contribution of SFR Santé Lyon-Est (UAR3453 CNRS, US7 Inserm, UCBL) facilitiy : CIQLE (a LyMIC member) for support and access to equipment, and Camilla Luccardini for technical assistance. Some strains were generated by SEGiCel (SFR Santé Lyon Est CNRS UAR 3453, Lyon, France). Some strains were provided by the Caenorhabditis Genetics Center (CGC), which is funded by the NIH Office of Research Infrastructure Programs (P40 OD010440). Funding for this study was provided by ANR Synapunct 580 ANR-22CE16-0024-01, ERC_Adg C.NAPSE #695295, Équipe FRM 2023 EQU202303016267, ANR-11-LABX-0042/ANR-11-IDEX-0007.

## Author contributions

Conceptualization: MC and JLB; Methodology: MC, LG, SLR, AD, AW and JLB; Investigation: MC, LG, SLR, AD, OR, CV, MJ, DLG and ACV; Formal analysis: MC, LG, SLR and AD; Visualization: MC; Writing – Original draft: MC and JLB; Writing – Review and editing: MC, LG, SLR, AD, MJ and JLB; Funding Acquisition: MC and JLB; Supervision: MC and JLB.

## Declaration of generative AI and AI-assisted technologies in the writing process

During the preparation of this work the authors used ChatGPT - OpenAI and DeepL in order to improve the readability and language of the manuscript. After using these tools, the authors reviewed and edited the content as needed and take full responsibility for the content of the publication.

## Material and methods

### Strains and genetics

All *C. elegans* strains were grown at 20°C on nematode growth medium (NGM) agar plates with *Escherichia coli* OP50 as a food source. All strains were originally derived from the wild-type Bristol N2 strain. A complete list of strains used in this study is provided in Table S1.

### EMS screen

Worms were collected and washed with M9 buffer (3 g of KH2PO4, 6 g of Na2HPO4, 5 g of NaCl and 0.25 g of MgSO4·7 H2O, distilled water up to 1 L) from 10 plates containing a majority of L4 (4^th^ larval stage). Mutagenesis was performed by incubating 4 mL of M9 containing the pellet of worms (P0) with 20 μL of EMS (SIGMA M-0880) for 4 h with agitation. Worms were then washed 5 times in 15 mL of M9 and put back on NGM plates with OP50. The next day, 40 P0 worms were transferred to individual plates to lay eggs (F1) and moved to fresh plates every day. When the progeny of F1 worms reached the adult stage, we analyzed their fluorescence using a Nikon AZ100 multizoom microscope and we cloned worms with an altered fluorescence pattern or level. After confirmation of the mutant phenotype in the next generation, we performed two rounds of crossing with the strain used for mutagenesis to remove most of the non-causal mutations induced by EMS. We then prepared genomic DNA (DNeasy blood and tissue kit, Qiagen) from pooling of 5 independent F2 from the 2^nd^ cross for each mutant strain and we sent it for whole genome sequencing (Novogene). Identification of EMS-induced mutations and *in silico* complementation were performed using in-house built scripts. In total, we performed 3 rounds of genetic screen. The first two rounds of screening were performed on a strain where there is a deletion of one nucleotide in the start codon of PunctinS (ΔG42), thus the tagRFP-T signal corresponds to PunctinL (PunctinL-RFP). These screens retrieved five mutations in the *cle-1* locus: C538Y, G575E, Q881*, G1062E and W1077* (Fig. 1B), the amino acid position corresponds to isoform A. The third round of screening was performed on a strain where the start codon of PunctinL isoforms was mutated from ATG to ATA (PunctinS-RFP). This screen retrieved two more alleles of *cle-1*: R450* and G470D (Fig. 1B).

### CRISPR/Cas9 genome engineering

All knock-in alleles were generated according to Ghanta and Mello 2020. crRNA was designed with the Benchling software and synthesized by Integrated DNA Technologies (IDT). A mix containing 2.8 μL crRNA (34 μM), 5 μL tracrRNA (18 μM, IDT #1073190) and 0.5 μL Cas9 nuclease (10 mL/mg, IDT #1081058) was incubated for 15 min at 37°C. For the generation of deletions, 2 crRNAs were used, at both ends of the deleted sequence. In this case, the mix contained 1.4 μL of each crRNA. A mix of 2.2 μL single strand repair template (1 μg/μL) or 500 ng double strand repair template, with 800 ng of pRF4 plasmid and molecular biology grade water up to 20 mL was added to the mix. This mix was injected in adult worms. Worms with a roller phenotype in the next generation were isolated and tested by PCR. All gene editions were confirmed by sequencing. Finally, the edited strains were outcrossed once. For *cle-1A/D* transcriptional reporter, incorrect insertion introduced a mutation in the first exon. A list of CRISPR alleles generated in this study and corresponding crRNAs is provided in Tables S2 and S3.

### Plasmid construction

The plasmids constructed for this study are described in Table S4. All constructs were verified by Sanger sequencing (Eurofins). Briefly, 1.5 μL plasmid was digested with 0.5 μL of one or two restriction enzymes (FastDigest, ThermoFisher Scientific) at 37°C. dsDNA fragments to integrate to the plasmid were generated by PCR and assembled with the digested plasmid using the Gibson cloning method^55^. Plasmids were then amplified in bacteria, purified with the PureLink™ HQ Mini Plasmid DNA Purification Kit (Invitrogen K210001) and verified using Sanger sequencing (Eurofins).

### Generation of single-copy insertion alleles

A list of single-copy alleles generated in this study is provided in Table S4. Single-copy insertion alleles were generated by the miniMos method^56^. For tissue-specific expression, the promoters used were as follow: *Punc-17* (cholinergic neurons) and *Punc-47* (GABAergic neurons). Worms were injected with 15 ng/μL plasmid of interest containing the promoter and the open reading frame fused to fluorescent proteins, 50 ng/μL pCFJ601 (Mos1 transposase), 10 ng/μL pMA122 (negative selective marker *Phsp16.2::peel-1*), and 2.5 ng/μL pCFJ90 (*Pmyo-2::mCherry*). Neomycin (G418) was added to plates 24 h after injection at 1.5 μg/μL final concentration. Candidate plates were heat shocked for 2 h at 34°C. Selected lines were then bred to homozygosity.

### Immunohistochemistry

Immunohistochemistry was performed as previously described^13^. In brief, *C. elegans* were freeze-cracked and fixed during 30 min in ice-cold paraformaldehyde (thermo scientific 28908, 4% in PBS) and washed 3 times in PBS. Blocking was performed by incubating worms in PBS supplemented with 0.2% fish gelatin (Sigma G7765) and 0.25% triton X-100 (Sigma T9284) for 1 h at room temperature. Primary antibodies were diluted in PGT solution (PBS, 0.1% fish gelatin, 0.25% triton X-100) and applied overnight at 4°C. Primary antibodies used here are anti-MYC (Genetex GTX29106, 1:5000) and anti-T7 (Novagen 69522-3, 1:500). Worms were then washed 5 times in PBS with 0.25% triton X-100. Worms were then incubated in secondary antibodies, diluted in PGT solution, for 2 h at room temperature. Secondary antibodies used here are the goat anti-mouse Alexa Fluor 647 (ThermoFisher scientific A-21235, 1:500) and the goat anti-rabbit Cy3 (ThermoFisher scientific A10520, 1:500). Worms were washed 5 times in PBS with 0.25% triton X-100 and mounted in Vectashield medium. Images were acquired using a spinning disk confocal microscope (see below).

### Live Confocal Microscopy and quantification

Animals were imaged at the young adult stage (24 h post L4 larval stage) unless specified otherwise. For i*n vivo* imaging, live hermaphrodites were mounted on 2% agarose (w/v in water) dry pads immersed in 5% poly-lysine beads (#00876-15 Biovalley) diluted in M9 buffer. Confocal images were taken on an Andor spinning disk system (Oxford Instruments) installed on a Nikon-IX86 microscope (Olympus) equipped with a 40X/NA1.3 and a 60X/NA1.4 oil immersion objective and an Evolve EMCCD camera. For mosaic imaging of entire adult worms, a grid of images covering the worm was acquired at 40X with a Z-stack. Images were then projected in Z (maximum intensity projection) and stitched in Fiji. For quantifications of fluorescence levels and colocalization, an image of the dorsal nerve cord (DNC) was acquired as a stack of optical sections (0.2-0.3 μm apart) for each animal. Images of the DNC were acquired around the mid-body, anterior to the vulva. Quantification of the signal intensity at the cord was performed using a macro in the Fiji software^57,58^. Briefly, images of DNCs were cropped as a rectangle containing solely the nerve cord and projected as a sum in Z direction. Then, the mean intensity was projected in x direction (orthogonal to the cord) and the intensity was measured as the area under the curve, after exclusion of the background. To quantify the colocalization between 2 channels, the 2D Pearson correlation coefficient was calculated on the summed projection using the JACoP plugin^59^ in Fiji. Plot profiles were generated in Fiji from the summed projection after rolling ball background subtraction and plotted in Prism 10 (GraphPad).

### Data representation and statistics

Confocal imaging of control and test groups were repeated at least 2 times for each experiment and the data were pooled. Fluorescence intensity was normalized to the mean value of the control group. Graphs were generated using the Prism 10 software (GraphPad): each dot represent a single worm (N numbers are indicated on each graph), histograms show the mean value and error bars correspond to the standard deviation. Statistical testing was performed using the Prism software and P values are indicated on each graph as follows: ns = non-significant, * p<0.05, ** p<0.01 and *** p<0.001. The normality of the distribution was assessed using the D’Agostino Pearson test. If the test returned a significant result for at least one genotype, a non-parametrical test was applied: the Mann-Whitney test for 2-group comparisons and the Kruskal-Wallis test with the Dunn’s multiple comparisons test for >2-group comparisons. If all genotypes have a normal distribution, variances were compared using the F test for 2-group comparisons and the Brown-Forsythe test for >2-group comparisons. If this test returned a non-significant result, groups were compared using the Student’s t test for 2-group comparisons and ordinary one-way ANOVAs with Tukey’s multiple comparisons test for >2-group comparisons. If variances significantly differed between genotypes, groups were compared using the Welch’s t test for 2-group comparisons and a Welch’s ANOVA with the Dunnett’s T3 multiple comparisons test for >2-group comparisons.

### Immunoprecipitations from worm lysates

Mixed-stage worms were collected from ten 10-cm dishes with 0.1 mM NaCl. After washing 3 times at 4°C, worm slurry was frozen in liquid nitrogen to form ∼5 mL of worm beads. Lysis was performed by manually grinding ∼2.5 mL of worm beads in liquid nitrogen and then incubating the resulting powder in 3.5 mL ice-cold RIPA buffer (50 mM Tris-HCl pH=7,5 ; 150 mM NaCl; 1% Triton, 0.1% SDS, 5mM EDTA pH=8; 1mM DTT; 0,5 % sodium deoxycholate) for 2 h at 4°C with rotation. This lysate was then centrifuged at 5,000 × *g* for 10 min at 4°C. Supernatant was collected and centrifuged at 16,000 × *g* for 15 min at 4°C. Supernatant was collected and diluted 1:1 in WRB1 buffer (50 mM Hepes, 50 mM KCl, 100 mM NaCl, 1 mM EDTA and one tablet of Roche-Merck cOmplete Protease inhibitor cocktail #05056489001 in 50 mL). Precleaned 20 μL Trap-A beads coupled with anti-GFP (Chromotek, gta), anti-RFP (Chromotek, rta) or anti-MYC (Chromotek, yta) were incubated overnight at 4°C with the samples. The beads were collected by centrifugation at 1500 × *g* at 4 °C and washed 3 times with WRB1 buffer. Precipitated proteins were eluted in 100 μL WRB1 and 40 μL Laemmi buffer 4X by heating at 95 °C for 10 min and centrifuged 1500 × g at RT for 5 min. 20 μL of supernatant was loaded to 4-20% (BioRad Mini-PROTEAN TGX Stain-Free Protein Gels, 4568094) or 8% (Clinisciences nUView Tris-Glycine Precast, NB10-008) acrylamide gels. Proteins were transferred to a Nitrocellulose membrane using the Trans-blot turbo transfer system (BIO-RAD) with the “mixed molecular weight” program. After 1 h of blocking in TBST (200 mM Tris; 1500 mM NaCl, 0.1 % Tween 20) and 5% low-fat milk (w/v in TBST), membranes were incubated overnight at 4°C with anti-GFP (1:2,000, Roche #11814460001), anti-RFP (1:1,000, Chromotek 6G6) or anti-MYC (1:1,000, Sigma M4439) primary antibodies. The membranes were then washed 3 times with TBST and incubated 30 min with HRP conjugated secondary anti-mouse antibody (1:3000, Cell signaling 7076S). Membranes were washed again 3 times with TBST, incubated 5 min in Clarity Max Western electrochemiluminescent substrate (BIO-RAD #175062) and using the Bio-rad ChemiDoc XRS system.

### Structure prediction

Protein structure prediction of PunctinL (AF-P90884-F1-v4) from AlphaFold Protein Structure Database^22–24^ was visualized using PyMOL Molecular Graphics System (Version 2.5.2).

### Data availability

All strains and reagents are available upon request. Table S1 provides a list of strains used in this study. Table S2 provides a list of alleles used in this study and Table S3 a list of the crRNAs used to generate the CRISPR alleles used in this study. Table S4 provides a list of plasmids and corresponding alleles generated in this study.

## Supplementary figure legends

**Supplementary Figure 1:**
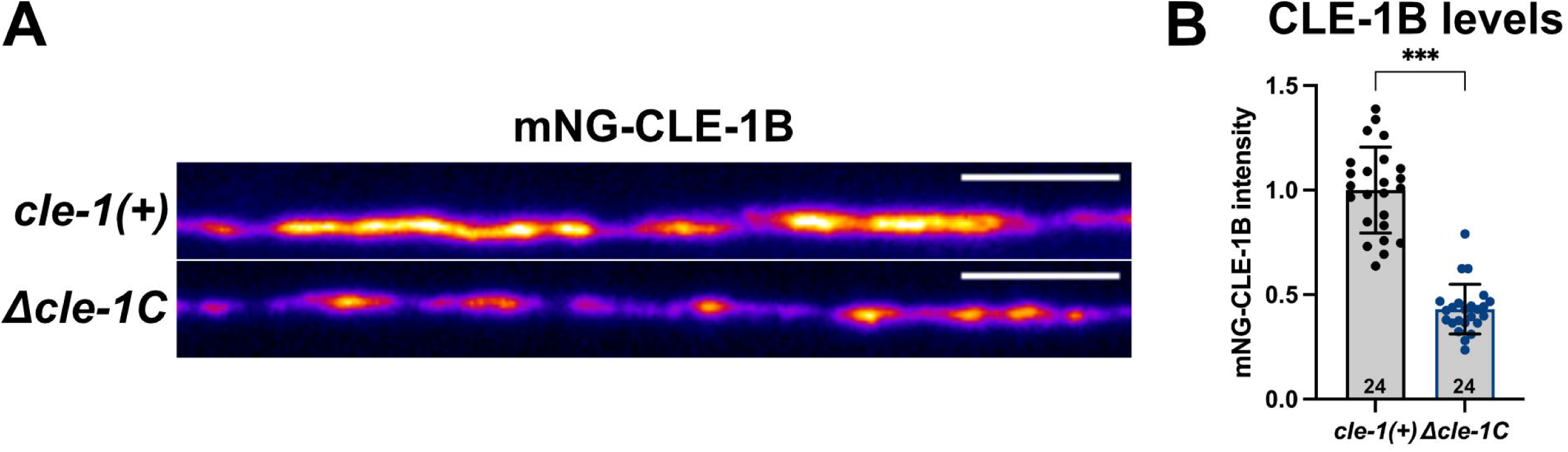
Levels of CLE-1B are reduced in *Δcle-1C* mutants. A, Confocal images of mNG-CLE-1B (*syb6520*) in controls and *Δcle-1C* mutants. Scale bar is 5 μm. B, Quantification of mNG-CLE-1B fluorescence levels at DNCs in controls and *cle-1(Q881*)* mutants (Mann-Whitney’s test).

**Supplementary Figure 2:**
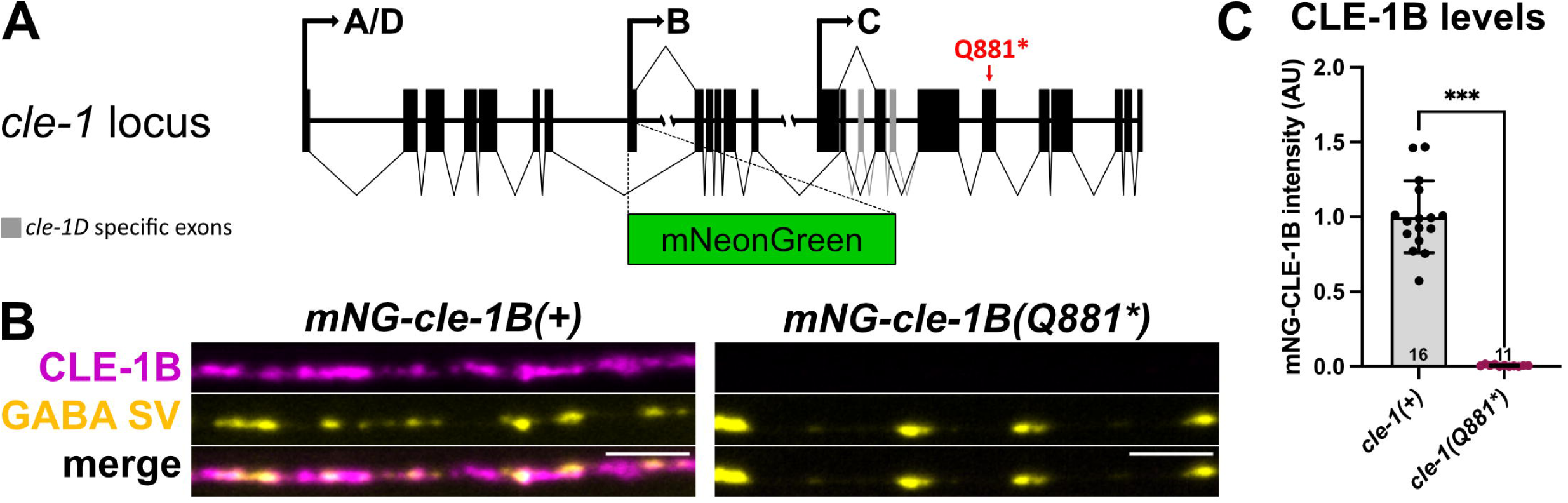
*Q881\** mutation abolishes the expression of CLE-1B. A, Schematic organization of the *cle-1* locus insertion of mNG and Q881* mutation in *mNG-cle-1B(Q881*)* mutants. B, Confocal images of mNG-CLE-1B (*syb6520*, magenta) and GABAergic SVs (*krIs67*, yellow) in controls and *cle-1(Q881*)* mutants. Scale bar is 5 μm. C, Quantification of mNG-CLE-1B fluorescence levels at DNCs in controls and *cle-1(Q881*)* mutants (Welch’s t test).

**Supplementary Figure 3:**
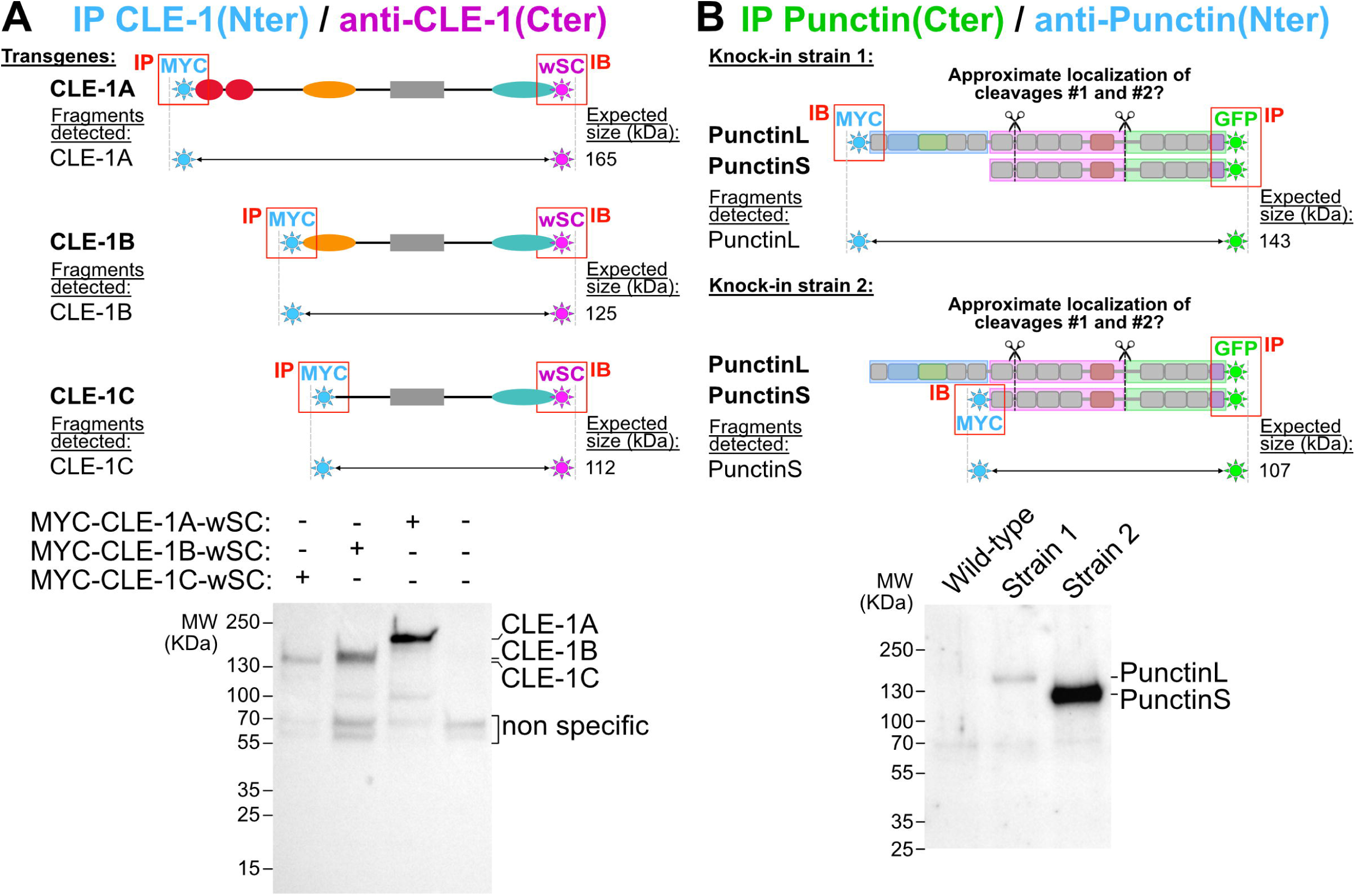
Identification of full-length CLE-1 and Punctin isoforms. A, Schematics of fusion proteins and immunoprecipitation of different CLE-1 isoforms (IP MYC) followed by anti-wSC blotting in wild types (lane 4), MYC-CLE1A-wSC (lane 1), MYC-CLE1B-wSC (lane 2) and MYC-CLE1C-wSC (lane 3). B, Schematics of fusion proteins and immunoprecipitation of Punctin(Cter) (IP GFP) followed by anti-MYC blotting in wild types (lane 1), MYC-PunctinL KI (lane 2, *kr421*, MYC-PunctinL-GFP+T7-PunctinS-GFP) and MYC-PunctinS KI (lane 3, *kr525*, PunctinL-GFP+MYC-PunctinS-GFP).

**Supplementary Figure 4:**
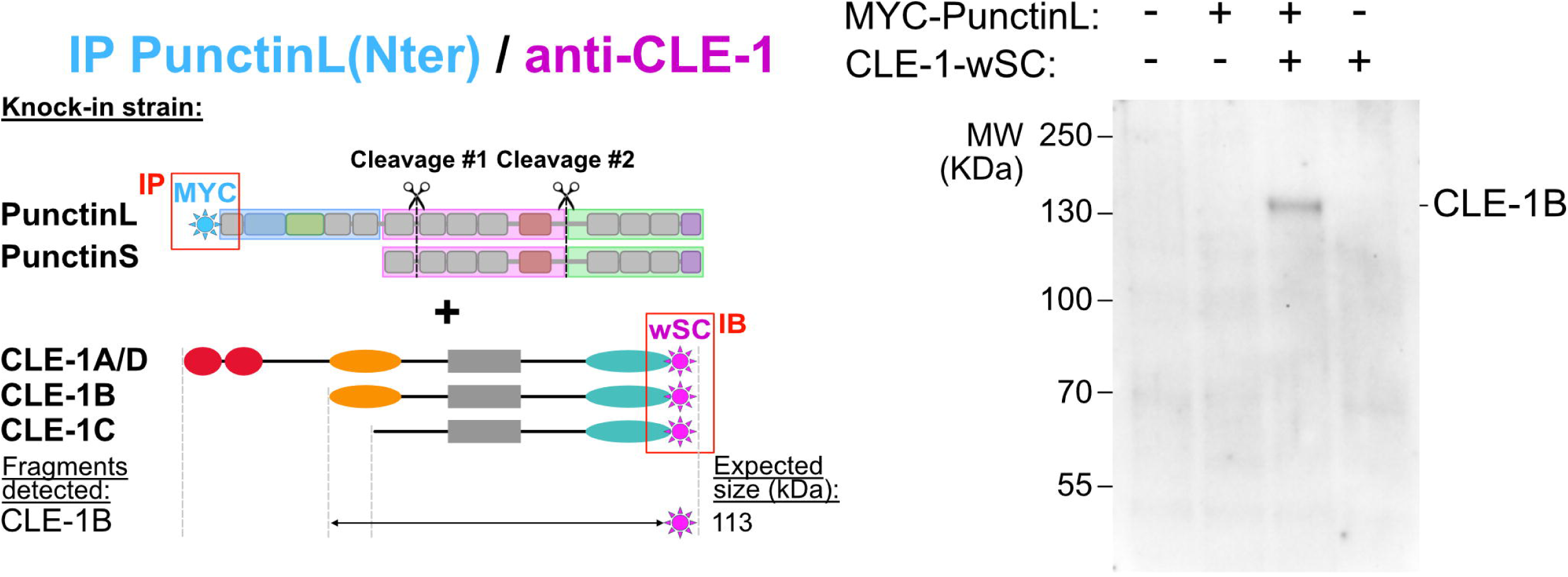
PunctinL forms a molecular complex with CLE-1B. Schematics of fusion proteins and immunoprecipitation of PunctinL(Nter) (IP MYC) followed by anti-wSC blotting of CLE-1 in wild types (lane 1), MYC-PunctinL KI (lane 2), MYC-PunctinL + CLE-1-wSC double KI (lane 3) and CLE-1-wSC KI (lane 4).

**Supplementary Figure 5:**
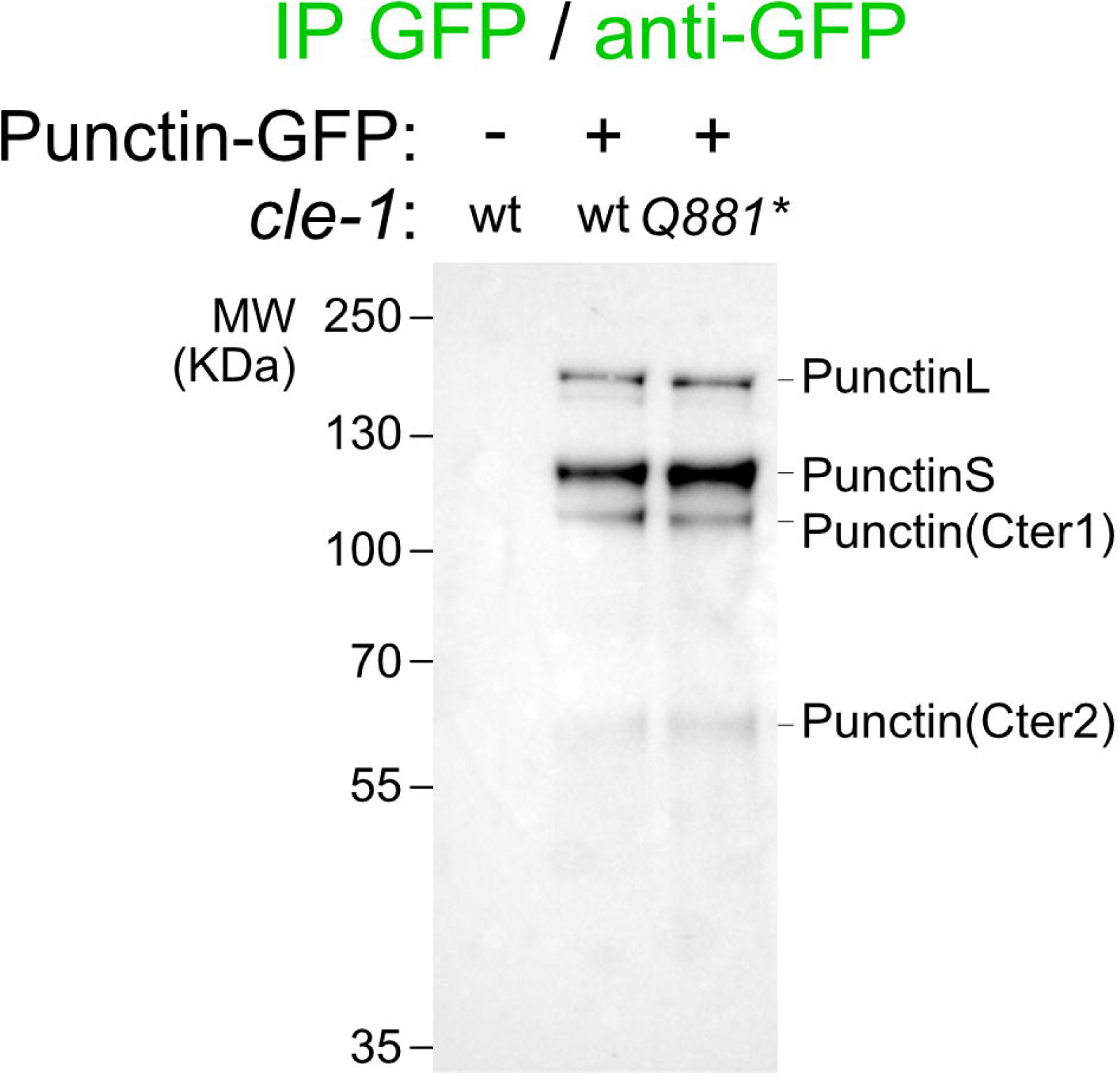
Punctin is correctly cleaved in *cle-1*mutants. Immunoprecipitation of Punctin-GFP followed by anti-GFP blotting in wild types (no tag, lane 1) and in Punctin-GFP KIs (strain 1) without (lane 2) or with *cle-1(Q881*)* mutation (lane 3).

**Supplementary Figure 6:**
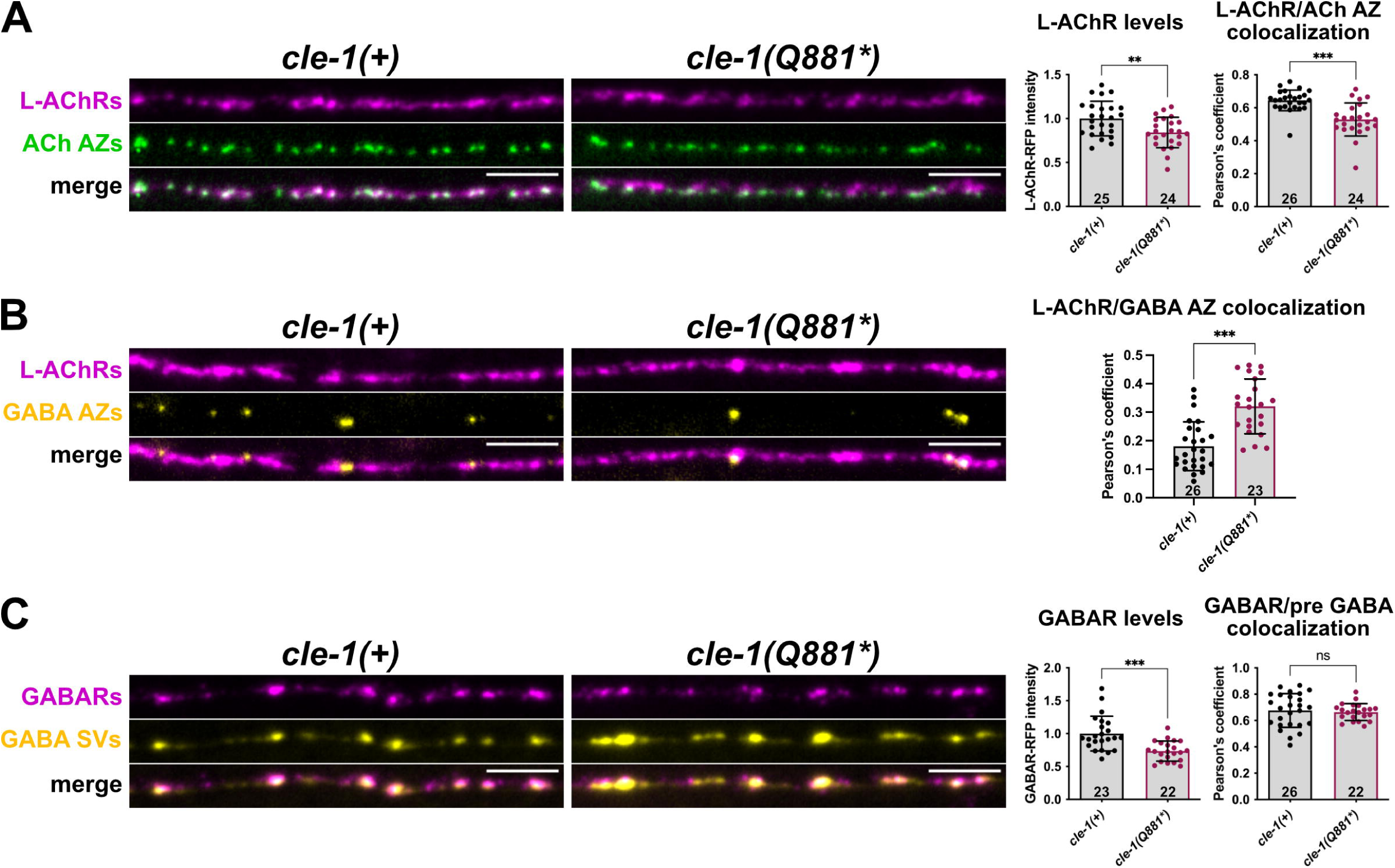
GABAR and L-AChR levels are mildly affected in *cle-1(Q881*)* mutants. A, Confocal images, quantifications of L-AChR-RFP intensity (*kr208*, magenta, Student’s t test) and colocalization between L-AChR-RFP and cholinergic AZs (*krSi145*, green, Mann-Whitney’s test) in controls and *cle-1(Q881*)* mutants. B, Confocal images and quantifications of the colocalization between L-AChR-RFP (*kr208*, magenta) and GABAergic AZs (*krSi141*, yellow, Student’s t test) in controls and *cle-1(Q881*)* mutants. C, Confocal images, quantifications of GABAR-RFP intensity (*kr296*, magenta, Mann-Whitney’s test) and colocalization between GABAR-RFP and GABA SVs (*krIs67*, yellow, Welch’s t test) in controls and *cle-1(Q881*)* mutants.

**Supplementary Figure 7:**
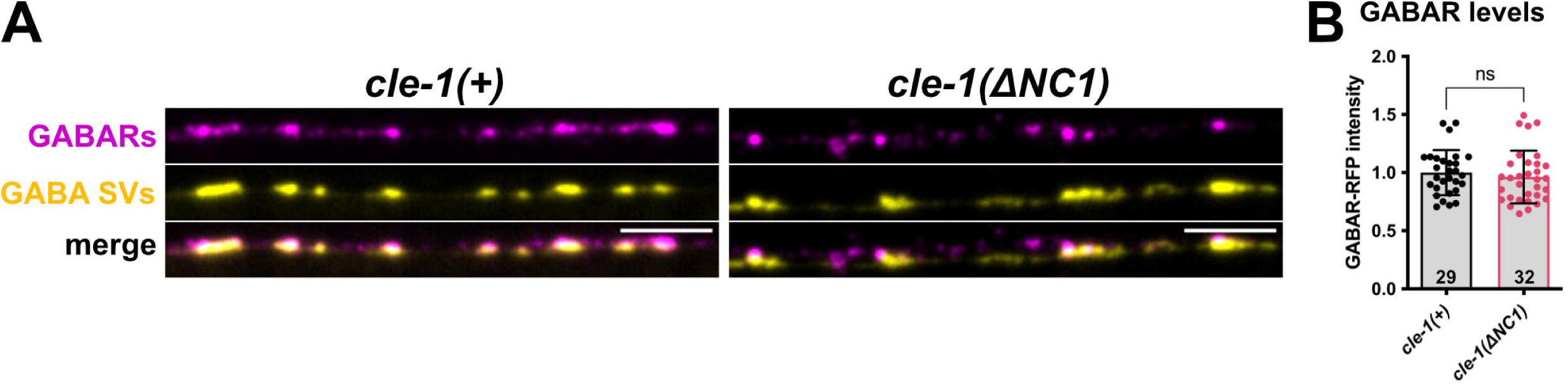
GABAR levels are unchanged in *cle-1(ΔNC1)* mutants. A, Confocal images of GABAR-RFP (*kr296*, magenta) and GABAergic SVs (*krIs67*, yellow) in controls and *cle-1(ΔNC1)* mutants. Scale bars are 5 μm. B, Quantifications of the GABAR-RFP fluorescence intensity at DNCs in controls and *cle-1(ΔNC1)* mutants. Student’s t test.

**Supplementary Figure 8:**
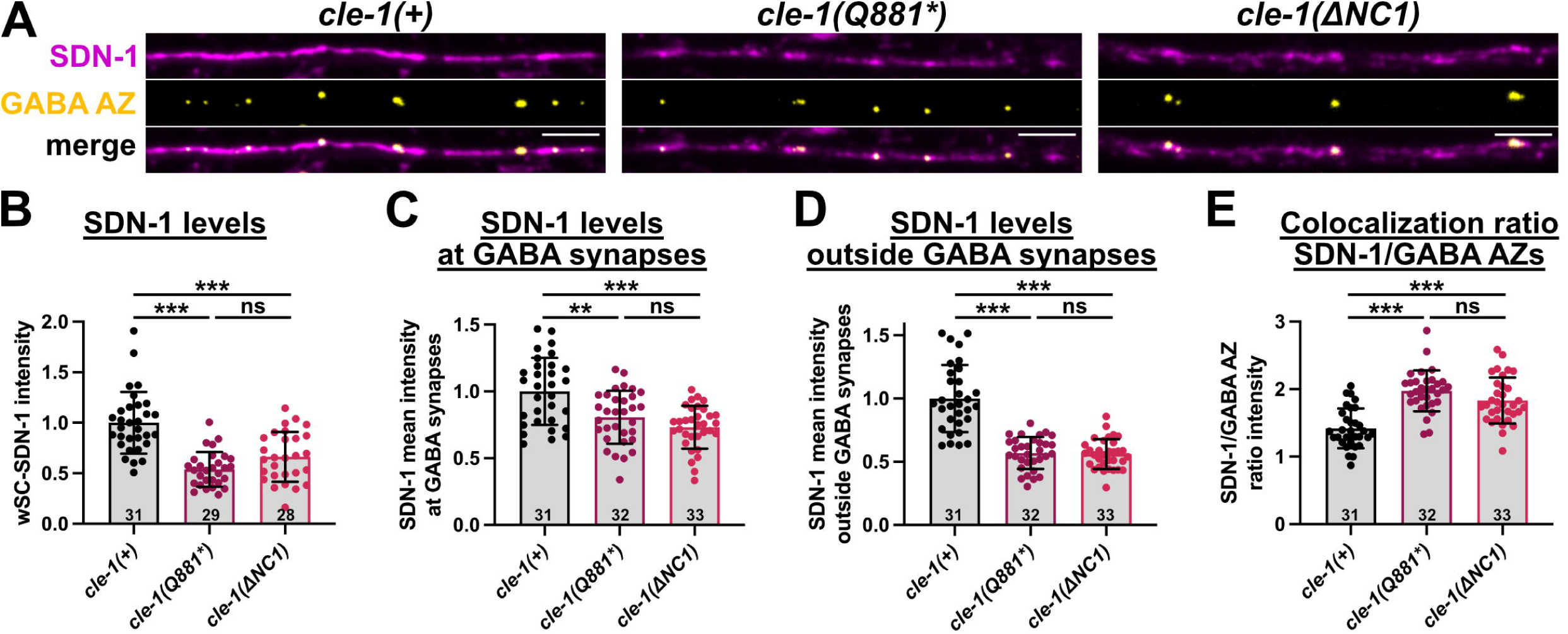
SDN-1/Syndecan is concentrated at GABAergic synapses in *cle-1* mutants. A, Confocal images of wSC-SDN-1 (*kr424*, magenta) and GABAergic AZs (*krSi141*, yellow) in controls, *cle-1(Q881*)* and *cle-1(ΔNC1)* mutants. B-E, Quantification of wSC-SDN-1 intensity at the cord (B, Kruskall-Wallis with Dunn’s multiple comparisons tests), at GABA synapses (C, Welch’s ANOVA with Dunett’s T3 multiple comparison tests), outside GABA synapses (D, Welch’s ANOVA with Dunett’s T3 multiple comparison tests) and colocalization ratio between the mean intensity of SDN-1 at GABA synapses and outside GABA synapses (E, One way ANOVA with Tukey’s multiple comparisons tests). Scale bars are 5 μm.

**Supplementary Figure 9:**
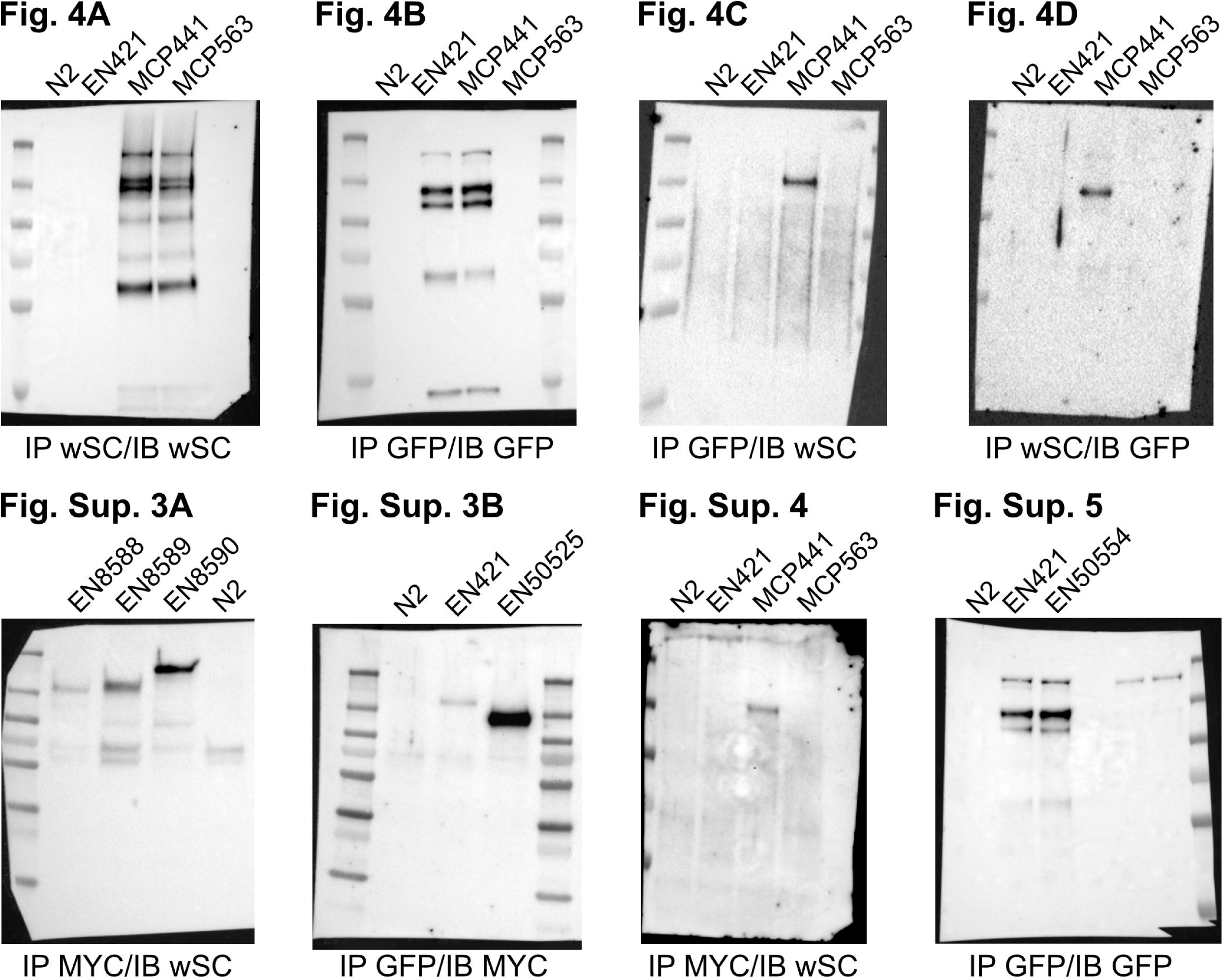
Full gel images. See corresponding figure for detailed legend.

**Supplementary Table 1: List of strains used in this study.**

**Supplementary Table 2: List of alleles used in this study.**

**Supplementary Table 3: List of crRNAs used in this study.**

**Supplementary Table 4: List of plasmids and corresponding alleles used in this study.**

## References

1. Südhof, T.C. (2018). Towards an Understanding of Synapse Formation. Neuron 100, 276–293. 10.1016/j.neuron.2018.09.040.

2. Yuzaki, M. (2018). Two Classes of Secreted Synaptic Organizers in the Central Nervous System. Annu. Rev. Physiol. 80, 243–262. 10.1146/annurev-physiol-021317-121322.

3. Matsuda, K., Miura, E., Miyazaki, T., Kakegawa, W., Emi, K., Narumi, S., Fukazawa, Y., Ito-Ishida, A., Kondo, T., Shigemoto, R., et al. (2010). Cbln1 Is a Ligand for an Orphan Glutamate Receptor δ2, a Bidirectional Synapse Organizer. Science 328, 363–368. 10.1126/science.1185152.

4. Uemura, T., Lee, S.-J., Yasumura, M., Takeuchi, T., Yoshida, T., Ra, M., Taguchi, R., Sakimura, K., and Mishina, M. (2010). Trans-Synaptic Interaction of GluRδ2 and Neurexin through Cbln1 Mediates Synapse Formation in the Cerebellum. Cell 141, 1068–1079. 10.1016/j.cell.2010.04.035.

5. Südhof, T.C. (2023). Cerebellin–neurexin complexes instructing synapse properties. Current Opinion in Neurobiology 81, 102727. 10.1016/j.conb.2023.102727.

6. Herbst, R., Huijbers, M.G., Oury, J., and Burden, S.J. (2024). Building, Breaking, and Repairing Neuromuscular Synapses. Cold Spring Harb Perspect Biol 16, a041490. 10.1101/cshperspect.a041490.

7. Irala, D., Wang, S., Sakers, K., Nagendren, L., Ulloa Severino, F.P., Bindu, D.S., Savage, J.T., and Eroglu, C. (2024). Astrocyte-secreted neurocan controls inhibitory synapse formation and function. Neuron 112, 1657–1675.e10. 10.1016/j.neuron.2024.03.007.

8. Christopherson, K.S., Ullian, E.M., Stokes, C.C.A., Mullowney, C.E., Hell, J.W., Agah, A., Lawler, J., Mosher, D.F., Bornstein, P., and Barres, B.A. (2005). Thrombospondins are astrocyte-secreted proteins that promote CNS synaptogenesis. Cell 120, 421–433. 10.1016/j.cell.2004.12.020.

9. Eroglu, C., Allen, N.J., Susman, M.W., O’Rourke, N.A., Park, C.Y., Ozkan, E., Chakraborty, C., Mulinyawe, S.B., Annis, D.S., Huberman, A.D., et al. (2009). Gabapentin receptor alpha2delta-1 is a neuronal thrombospondin receptor responsible for excitatory CNS synaptogenesis. Cell 139, 380–392. 10.1016/j.cell.2009.09.025.

10. González-Calvo, I., Cizeron, M., Bessereau, J.-L., and Selimi, F. (2022). Synapse Formation and Function Across Species: Ancient Roles for CCP, CUB, and TSP-1 Structural Domains. Front. Neurosci. 16, 866444. 10.3389/fnins.2022.866444.

11. Mizumoto, K., Jin, Y., and Bessereau, J.-L. (2023). Synaptogenesis: unmasking molecular mechanisms using *Caenorhabditis elegans*. GENETICS 223, iyac176. 10.1093/genetics/iyac176.

12. Cramer, T.M.L., Pinan-Lucarre, B., Cavaccini, A., Damilou, A., Tsai, Y.-C., Bhat, M.A., Panzanelli, P., Rama, N., Mehlen, P., Benke, D., et al. (2023). Adamtsl3 mediates DCC signaling to selectively promote GABAergic synapse function. Cell Reports 42, 112947. 10.1016/j.celrep.2023.112947.

13. Pinan-Lucarré, B., Tu, H., Pierron, M., Cruceyra, P.I., Zhan, H., Stigloher, C., Richmond, J.E., and Bessereau, J.-L. (2014). C. elegans Punctin specifies cholinergic versus GABAergic identity of postsynaptic domains. Nature 511, 466–470. 10.1038/nature13313.

14. Maro, G.S., Gao, S., Olechwier, A.M., Hung, W.L., Liu, M., Özkan, E., Zhen, M., and Shen, K. (2015). MADD-4/Punctin and Neurexin Organize C. elegans GABAergic Postsynapses through Neuroligin. Neuron 86, 1420–1432. 10.1016/j.neuron.2015.05.015.

15. Tu, H., Pinan-Lucarré, B., Ji, T., Jospin, M., and Bessereau, J.-L. (2015). C. elegans Punctin Clusters GABAA Receptors via Neuroligin Binding and UNC-40/DCC Recruitment. Neuron 86, 1407–1419. 10.1016/j.neuron.2015.05.013.

16. Zhou, X., Gueydan, M., Jospin, M., Ji, T., Valfort, A., Pinan-Lucarré, B., and Bessereau, J.-L. (2020). The netrin receptor UNC-40/DCC assembles a postsynaptic scaffold and sets the synaptic content of GABAA receptors. Nat Commun 11, 2674. 10.1038/s41467-020-16473-5.

17. Zhou, X., Vachon, C., Cizeron, M., Romatif, O., Bülow, H.E., Jospin, M., and Bessereau, J.-L. (2021). The HSPG syndecan is a core organizer of cholinergic synapses. Journal of Cell Biology 220, e202011144. 10.1083/jcb.202011144.

18. Ricard-Blum, S., and Vallet, S.D. (2016). Proteases decode the extracellular matrix cryptome. Biochimie 122, 300–313. 10.1016/j.biochi.2015.09.016.

19. Heljasvaara, R., Aikio, M., Ruotsalainen, H., and Pihlajaniemi, T. (2017). Collagen XVIII in tissue homeostasis and dysregulation — Lessons learned from model organisms and human patients. Matrix Biology 57–58, 55–75. 10.1016/j.matbio.2016.10.002.

20. Keeley, D.P., Hastie, E., Jayadev, R., Kelley, L.C., Chi, Q., Payne, S.G., Jeger, J.L., Hoffman, B.D., and Sherwood, D.R. (2020). Comprehensive Endogenous Tagging of Basement Membrane Components Reveals Dynamic Movement within the Matrix Scaffolding. Developmental Cell 54, 60–74.e7. 10.1016/j.devcel.2020.05.022.

21. Ackley, B.D., Crew, J.R., Elamaa, H., Pihlajaniemi, T., Kuo, C.J., and Kramer, J.M. (2001). The NC1/endostatin domain of Caenorhabditis elegans type XVIII collagen affects cell migration and axon guidance. J Cell Biol 152, 1219–1232. 10.1083/jcb.152.6.1219.

22. Jumper, J., Evans, R., Pritzel, A., Green, T., Figurnov, M., Ronneberger, O., Tunyasuvunakool, K., Bates, R., Žídek, A., Potapenko, A., et al. (2021). Highly accurate protein structure prediction with AlphaFold. Nature 596, 583–589. 10.1038/s41586-021-03819-2.

23. Varadi, M., Anyango, S., Deshpande, M., Nair, S., Natassia, C., Yordanova, G., Yuan, D., Stroe, O., Wood, G., Laydon, A., et al. (2022). AlphaFold Protein Structure Database: massively expanding the structural coverage of protein-sequence space with high-accuracy models. Nucleic Acids Research 50, D439–D444. 10.1093/nar/gkab1061.

24. Varadi, M., Bertoni, D., Magana, P., Paramval, U., Pidruchna, I., Radhakrishnan, M., Tsenkov, M., Nair, S., Mirdita, M., Yeo, J., et al. (2024). AlphaFold Protein Structure Database in 2024: providing structure coverage for over 214 million protein sequences. Nucleic Acids Research 52, D368–D375. 10.1093/nar/gkad1011.

25. Correa, E., Mialon, M., Cizeron, M., Bessereau, J.-L., Pinan-Lucarre, B., and Kratsios, P. (2024). UNC-30/PITX coordinates neurotransmitter identity with postsynaptic GABA receptor clustering. Development 151, dev202733. 10.1242/dev.202733.

26. Drechsler, M., Schmidt, A.C., Meyer, H., and Paululat, A. (2013). The Conserved ADAMTS-like Protein Lonely heart Mediates Matrix Formation and Cardiac Tissue Integrity. PLoS Genet 9, e1003616. 10.1371/journal.pgen.1003616.

27. Ackley, B.D., Kang, S.H., Crew, J.R., Suh, C., Jin, Y., and Kramer, J.M. (2003). The basement membrane components nidogen and type XVIII collagen regulate organization of neuromuscular junctions in Caenorhabditis elegans. J Neurosci 23, 3577–3587. 10.1523/JNEUROSCI.23-09-03577.2003.

28. Qin, J., Liang, J., and Ding, M. (2014). Perlecan Antagonizes Collagen IV and ADAMTS9/GON-1 in Restricting the Growth of Presynaptic Boutons. Journal of Neuroscience 34, 10311–10324. 10.1523/JNEUROSCI.5128-13.2014.

29. Lemons L, M., McKillop, H., Genao, N., and Francis M, M. (2024). The extracellular matrix molecule Collagen XVIII/CLE-1 affects neuronal dendritic spines. MicroPubl Biol 2024. 10.17912/micropub.biology.001331.

30. Bretaud, S., Guillon, E., Karppinen, S.-M., Pihlajaniemi, T., and Ruggiero, F. (2020). Collagen XV, a multifaceted multiplexin present across tissues and species. Matrix Biology Plus 6–7, 100023. 10.1016/j.mbplus.2020.100023.

31. Cunningham, C., Bolcaen, J., Bisio, A., Genis, A., Strijdom, H., and Vandevoorde, C. (2023). Recombinant Endostatin as a Potential Radiosensitizer in the Treatment of Non-Small Cell Lung Cancer. Pharmaceuticals 16, 219. 10.3390/ph16020219.

32. Méndez-Valdés, G., Gómez-Hevia, F., Lillo-Moya, J., González-Fernández, T., Abelli, J., Cereceda-Cornejo, A., Bragato, M.C., Saso, L., and Rodrigo, R. (2023). Endostatin and Cancer Therapy: A Novel Potential Alternative to Anti-VEGF Monoclonal Antibodies. Biomedicines 11, 718. 10.3390/biomedicines11030718.

33. Kuo, C.J., LaMontagne, K.R., Garcia-Cardeña, G., Ackley, B.D., Kalman, D., Park, S., Christofferson, R., Kamihara, J., Ding, Y.-H., Lo, K.-M., et al. (2001). Oligomerization-Dependent Regulation of Motility and Morphogenesis by the Collagen Xviii Nc1/Endostatin Domain. The Journal of Cell Biology 152, 1233–1246. 10.1083/jcb.152.6.1233.

34. Wang, T., Hauswirth, A.G., Tong, A., Dickman, D.K., and Davis, G.W. (2014). Endostatin Is a Trans-Synaptic Signal for Homeostatic Synaptic Plasticity. Neuron 83, 616–629. 10.1016/j.neuron.2014.07.003.

35. Wang, T., Morency, D.T., Harris, N., and Davis, G.W. (2020). Epigenetic Signaling in Glia Controls Presynaptic Homeostatic Plasticity. Neuron 105, 491–505.e3. 10.1016/j.neuron.2019.10.041.

36. Su, J., Stenbjorn, R.S., Gorse, K., Su, K., Hauser, K.F., Ricard-Blum, S., Pihlajaniemi, T., and Fox, M.A. (2012). Target-Derived Matricryptins Organize Cerebellar Synapse Formation through α3β1 Integrins. Cell Reports 2, 223–230. 10.1016/j.celrep.2012.07.001.

37. Wei, P., Pattarini, R., Rong, Y., Guo, H., Bansal, P.K., Kusnoor, S.V., Deutch, A.Y., Parris, J., and Morgan, J.I. (2012). The Cbln family of proteins interact with multiple signaling pathways. Journal of Neurochemistry 121, 717–729. 10.1111/j.1471-4159.2012.07648.x.

38. Haddick, P.C.G., Tom, I., Luis, E., Quiñones, G., Wranik, B.J., Ramani, S.R., Stephan, J.-P., Tessier-Lavigne, M., and Gonzalez, L.C. (2014). Defining the Ligand Specificity of the Deleted in Colorectal Cancer (DCC) Receptor. PLoS ONE 9, e84823. 10.1371/journal.pone.0084823.

39. Rinta-Jaskari, M.M., Naillat, F., Ruotsalainen, H.J., Koivunen, J.T., Sasaki, T., Pietilä, I., Elamaa, H.P., Kaur, I., Manninen, A., Vainio, S.J., et al. (2023). Temporally and spatially regulated collagen XVIII isoforms are involved in ureteric tree development via the TSP1-like domain. Matrix Biology 115, 139–159. 10.1016/j.matbio.2023.01.001.

40. Munezane, H., Oizumi, H., Wakabayashi, T., Nishio, S., Hirasawa, T., Sato, T., Harada, A., Yoshida, T., Eguchi, T., Yamanashi, Y., et al. (2019). Roles of Collagen XXV and Its Putative Receptors PTPσ/δ in Intramuscular Motor Innervation and Congenital Cranial Dysinnervation Disorder. Cell Reports 29, 4362–4376.e6. 10.1016/j.celrep.2019.11.112.

41. Ricard-Blum, S., and Salza, R. (2014). Matricryptins and matrikines: biologically active fragments of the extracellular matrix. Experimental Dermatology 23, 457–463. 10.1111/exd.12435.

42. Hirohata, S., Wang, L.W., Miyagi, M., Yan, L., Seldin, M.F., Keene, D.R., Crabb, J.W., and Apte, S.S. (2002). Punctin, a Novel ADAMTS-like Molecule, ADAMTSL-1, in Extracellular Matrix. Journal of Biological Chemistry 277, 12182–12189. 10.1074/jbc.M109665200.

43. Hall, N.G., Klenotic, P., Anand-Apte, B., and Apte, S.S. (2003). ADAMTSL-3/punctin-2, a novel glycoprotein in extracellular matrix related to the ADAMTS family of metalloproteases. Matrix Biology 22, 501–510. 10.1016/S0945-053X(03)00075-1.

44. Taye, N., Redhead, C., and Hubmacher, D. (2024). Secreted ADAMTS-like proteins as regulators of connective tissue function. American Journal of Physiology-Cell Physiology 326, C756–C767. 10.1152/ajpcell.00680.2023.

45. Dow, D.J., Huxley-Jones, J., Hall, J.M., Francks, C., Maycox, P.R., Kew, J.N.C., Gloger, I.S., Mehta, N.A.L., Kelly, F.M., Muglia, P., et al. (2011). ADAMTSL3 as a candidate gene for schizophrenia: Gene sequencing and ultra-high density association analysis by imputation. Schizophrenia Research 127, 28–34. 10.1016/j.schres.2010.12.009.

46. Irene Díez García-Prieto, I., Lopez-Martín, S., Albert, J., Jiménez de la Peña, M., Fernández-Mayoralas, D.M., Calleja-Pérez, B., Gómez Fernández, M.T., Álvarez, S., Pihlajaniemi, T., Izzi, V., et al. (2022). Mutations in the COL18A1 gen associated with knobloch syndrome and structural brain anomalies: a novel case report and literature review of neuroimaging findings. Neurocase 28, 11–18. 10.1080/13554794.2021.1928228.

47. Suzuki, O.T., Sertié, A.L., Der Kaloustian, V.M., Kok, F., Carpenter, M., Murray, J., Czeizel, A.E., Kliemann, S.E., Rosemberg, S., Monteiro, M., et al. (2002). Molecular analysis of collagen XVIII reveals novel mutations, presence of a third isoform, and possible genetic heterogeneity in Knobloch syndrome. Am J Hum Genet 71, 1320– 1329. 10.1086/344695.

48. Corbett, M.A., Turner, S.J., Gardner, A., Silver, J., Stankovich, J., Leventer, R.J., Derry, C.P., Carroll, R., Ha, T., Scheffer, I.E., et al. (2017). Familial epilepsy with anterior polymicrogyria as a presentation of COL18A1 mutations. European Journal of Medical Genetics 60, 437–443. 10.1016/j.ejmg.2017.06.002.

49. Deininger, M.H., Fimmen, B.A., Thal, D.R., Schluesener, H.J., and Meyermann, R. (2002). Aberrant Neuronal and Paracellular Deposition of Endostatin in Brains of Patients with Alzheimer’s Disease. J. Neurosci. 22, 10621–10626. 10.1523/JNEUROSCI.22-24-10621.2002.

50. van Horssen, J., Wilhelmus, M.M.M., Heljasvaara, R., Pihlajaniemi, T., Wesseling, P., de Waal, R.M.W., and Verbeek, M.M. (2002). Collagen XVIII: a novel heparan sulfate proteoglycan associated with vascular amyloid depositions and senile plaques in Alzheimer’s disease brains. Brain Pathol 12, 456–462. 10.1111/j.1750-3639.2002.tb00462.x.

51. Hendee, K., Wang, L.W., Reis, L.M., Rice, G.M., Apte, S.S., and Semina, E.V. (2017). Identification and functional analysis of an ADAMTSL1 variant associated with a complex phenotype including congenital glaucoma, craniofacial, and other systemic features in a three-generation human pedigree. Hum Mutat 38, 1485–1490. 10.1002/humu.23299.

52. Liu, Y., Zhang, J.-J., Piao, S.-Y., Shen, R.-J., Ma, Y., Xue, Z.-Q., Zhang, W., Liu, J., Jin, Z.-B., and Zhuang, W.-J. (2021). Whole-Exome Sequencing in a Cohort of High Myopia Patients in Northwest China. Front Cell Dev Biol 9, 645501. 10.3389/fcell.2021.645501.

53. Rudzinski, M.N., Chen, L., and Hernandez, M.R. (2008). Antiangiogenic characteristics of astrocytes from optic nerve heads with primary open-angle glaucoma. Arch Ophthalmol 126, 679–685. 10.1001/archopht.126.5.679.

54. Ghanta, K.S., and Mello, C.C. (2020). Melting dsDNA Donor Molecules Greatly Improves Precision Genome Editing in *Caenorhabditis elegans*. Genetics 216, 643– 650. 10.1534/genetics.120.303564.

55. Gibson, D.G., Young, L., Chuang, R.-Y., Venter, J.C., Hutchison, C.A., and Smith, H.O. (2009). Enzymatic assembly of DNA molecules up to several hundred kilobases. Nat Methods 6, 343–345. 10.1038/nmeth.1318.

56. Frøkjær-Jensen, C., Davis, M.W., Sarov, M., Taylor, J., Flibotte, S., LaBella, M., Pozniakovsky, A., Moerman, D.G., and Jorgensen, E.M. (2014). Random and targeted transgene insertion in Caenorhabditis elegans using a modified Mos1 transposon. Nat Methods 11, 529–534. 10.1038/nmeth.2889.

57. Schindelin, J., Arganda-Carreras, I., Frise, E., Kaynig, V., Longair, M., Pietzsch, T., Preibisch, S., Rueden, C., Saalfeld, S., Schmid, B., et al. (2012). Fiji: an open-source platform for biological-image analysis. Nat Methods 9, 676–682. 10.1038/nmeth.2019.

58. Rueden, C.T., Schindelin, J., Hiner, M.C., DeZonia, B.E., Walter, A.E., Arena, E.T., and Eliceiri, K.W. (2017). ImageJ2: ImageJ for the next generation of scientific image data. BMC Bioinformatics 18, 529. 10.1186/s12859-017-1934-z.

59. Bolte, S., and Cordelières, F.P. (2006). A guided tour into subcellular colocalization analysis in light microscopy. Journal of Microscopy 224, 213–232. 10.1111/j.1365-2818.2006.01706.x.

